# Linking carbohydrate structure with function in the human gut microbiome using hybrid metagenome assemblies

**DOI:** 10.1101/2021.05.11.441322

**Authors:** Anuradha Ravi, Perla Troncoso-Rey, Jennifer Ahn-Jarvis, Kendall R. Corbin, Suzanne Harris, Hannah Harris, Alp Aydin, Gemma L. Kay, Thanh Le Viet, Rachel Gilroy, Mark J. Pallen, Andrew J. Page, Justin O’Grady, Frederick J. Warren

**Author notes:** Contributed equally (Anuradha Ravi, Perla Troncoso-Rey, Jennifer Ahn-Jarvis, Justin O’Grady, Frederick J. Warren).

## Abstract

**Background:** Complex carbohydrates that escape digestion in the small intestine, are broken down in the large intestine by enzymes encoded by the gut microbiome. This is a symbiotic relationship between particular microbes and the host, resulting in metabolic products that influence host gut health and are exploited by other microbes. However, the role of carbohydrate structure in directing microbiota community composition and the succession of carbohydrate-degrading microbes is not fully understood. Here we take the approach of combining data from long and short read sequencing allowing recovery of large numbers of high quality genomes, from which we can predict carbohydrate degrading functions, and impact of carbohydrate on microbial communities.

**Results:** In this study we evaluate species-level compositional variation within a single microbiome in response to six structurally distinct carbohydrates in a controlled model gut using hybrid metagenome assemblies. We identified 509 high-quality metagenome-assembled genomes (MAGs) belonging to ten bacterial classes and 28 bacterial families. We found dynamic variations in the microbiome amongst carbohydrate treatments, and over time. Using these data, the MAGs were characterised as primary (0h to 6h) and secondary degraders (12h to 24h). Annotating the MAG’s with the Carbohydrate Active Enzyme (CAZyme) database we are able to identify species which are enriched through time and have the potential to actively degrade carbohydrate substrates.

**Conclusions:** Recent advances in sequencing technology allowed us to identify significant unexplored diversity amongst starch degrading species in the human gut microbiota including CAZyme profiles and complete MAGs. We have identified changes in microbial community composition in response to structurally distinct carbohydrate substrates, which can be directly related to the CAZyme complement of the enriched MAG’s. Through this approach, we have identified a number of species which have not previously been implicated in starch degradation, but which have the potential to play an important role.

---

Microbial diversity within the microbiome and its interactions with host health and nutrition are now widely studied[1]. An important role of the human gut microbiome is the metabolic breakdown of complex carbohydrates derived from plants and animals (e.g. legumes, seeds, tissue and cartilage)[2]. Short chain fatty acids (SCFA) are the main products of carbohydrate fermentation by gut microbiota and provide a myriad of health benefits through their systemic effects on host metabolism.[3, 4] However, we still do not have a complete picture of the range of microbial species involved in fermentation of complex carbohydrates to produce SCFA. Understanding the intricacies of complex carbohydrate metabolism by the gut microbiota is a significant challenge. The function of many ‘hard to culture species’ remains obscure and while advances in sequencing technology are beginning to reveal the true diversity of the human gut microbiota, there is still much to be learned.[5]

A key challenge is understanding the influence of structural complexity of carbohydrates on microbiota composition. Carbohydrates possess immense structural diversity, both at the chemical composition level (monomer and sugar linkage composition) and at the mesoscale. Individual species, or groups of species, within the gut microbiota are highly adapted to defined carbohydrate structures[6]. Starch is representative of the structural diversity found amongst carbohydrates and serves as a good model system as starches are readily fermented by several different species of colonic bacteria.[7] The gut microbiota is repeatedly presented with starches of diverse structures from the diet.[8] Consistent in starch is an α-1→4 linked glucose back bone, interspersed with α-1→6 linked branch points. Despite this apparent structural simplicity, starches botanical origin and subsequent processing (e.g. cooking) impacts its physicochemical properties, particularly crystallinity and recalcitrance to digestion.[7] It has been shown *in vitro*[7], in animal models[9] and in human interventions[8], that altering starch structure can have a profound impact on gut microbiome composition.

The microbiome is known to harbour a huge repertoire of carbohydrate-active enzymes (CAZymes) that can degrade diverse carbohydrate structures.[10, 11] However, it is a formidable challenge to study this functionality in complex microbial communities due to limitations in the depth of sequencing and coverage of all members in the community. While metagenomic sequencing has become a key tool, identifying genomes and functional pathways within the microbiome remains challenging in second generation sequencing due to limitations associated with short (∼300bp) reads. Third generation sequencing such as nanopore sequencing (Oxford Nanopore Technologies (ONT)) promises to circumvent these difficulties by providing longer reads (> 3 kilobase pairs [kbp]). This technology has become popular in clinical metagenomics for rapid pathogen diagnosis[12] and in human genomics research.[13] Long-read sequences can help bridge inter-genomic repeats and produce better de novo assembled genomes.[14] While the MinION platform from ONT has been used for metagenomic studies,[15] it cannot provide sufficient sequencing depth and coverage to sequence the many hundreds of genomes present in the human gut microbiome. PromethION (ONT) is capable of producing far greater numbers of sequences compared to either MinION or GridION, averaging four-five times more data per flow cell and the capacity to run up to 48 flow cells in parallel; this makes it suitable for metagenomics and microbiome studies. For example, PromethION has been used for long-read sequencing of environmental samples such as wastewater sludge, demonstrating its potential to recover large numbers of metagenome-assembled genomes (MAGs) from diverse mixtures of microbial species.[16] However, long error-prone reads aren’t ideal for species resolution metagenomics, therefore, a hybrid approach using short and long read data has been found to be most effective for generating accurate MAGs.[17]

To achieve species-level resolution of the microbes present in the gut microbiome during complex carbohydrate utilisation, we conducted a genome-resolved metagenomics study in a controlled gut colon model. *In vitro* fermentation systems have been used extensively to model changes in the gut microbial community as a result of external inputs, e.g., changes in pH, protein and carbohydrate supply[7, 18, 19]. We measured the dynamic changes in bacterial populations during fermentation of six structurally contrasting substrates: two highly recalcitrant starches (native Hylon VII (“Hylon”) and native potato starch (“potato”)); two accessible starches (native normal maize starch (“n.maize”) and gelatinized then retrograded maize starch (“r.maize”); an insoluble fibre (cellulose) resistant to fermentation (“Avicel”); and a highly fermentable soluble fibre (“inulin”). By generating hybrid assemblies using PromethION and NovaSeq data, we obtained 509 MAGs. The dereplicated set consisted of 151 genomes belonging to ten bacterial classes and 28 bacterial families. Using genome-level information and read proportions data, we identified several species that have novel putative starch-degrading properties.

## Results

### PromethION and NovaSeq sequencing of model gut samples enriched for carbohydrate degrading species

Fermentation of six contrasting carbohydrate substrates (inulin, Hylon, n.maize, potato, r.maize and Avicel; see methods section) was initiated by inoculation of the model colon with a carbohydrate and faecal material and the gut microbial community composition was monitored over time (0h, 6h, 12h and 24h) by sequencing as shown in Figure 1. In total, 23 samples and a negative control were sequenced (see Supplementary Table 1 for the PromethION and NovaSeq summary sequencing statistics).

**Figure 1.**
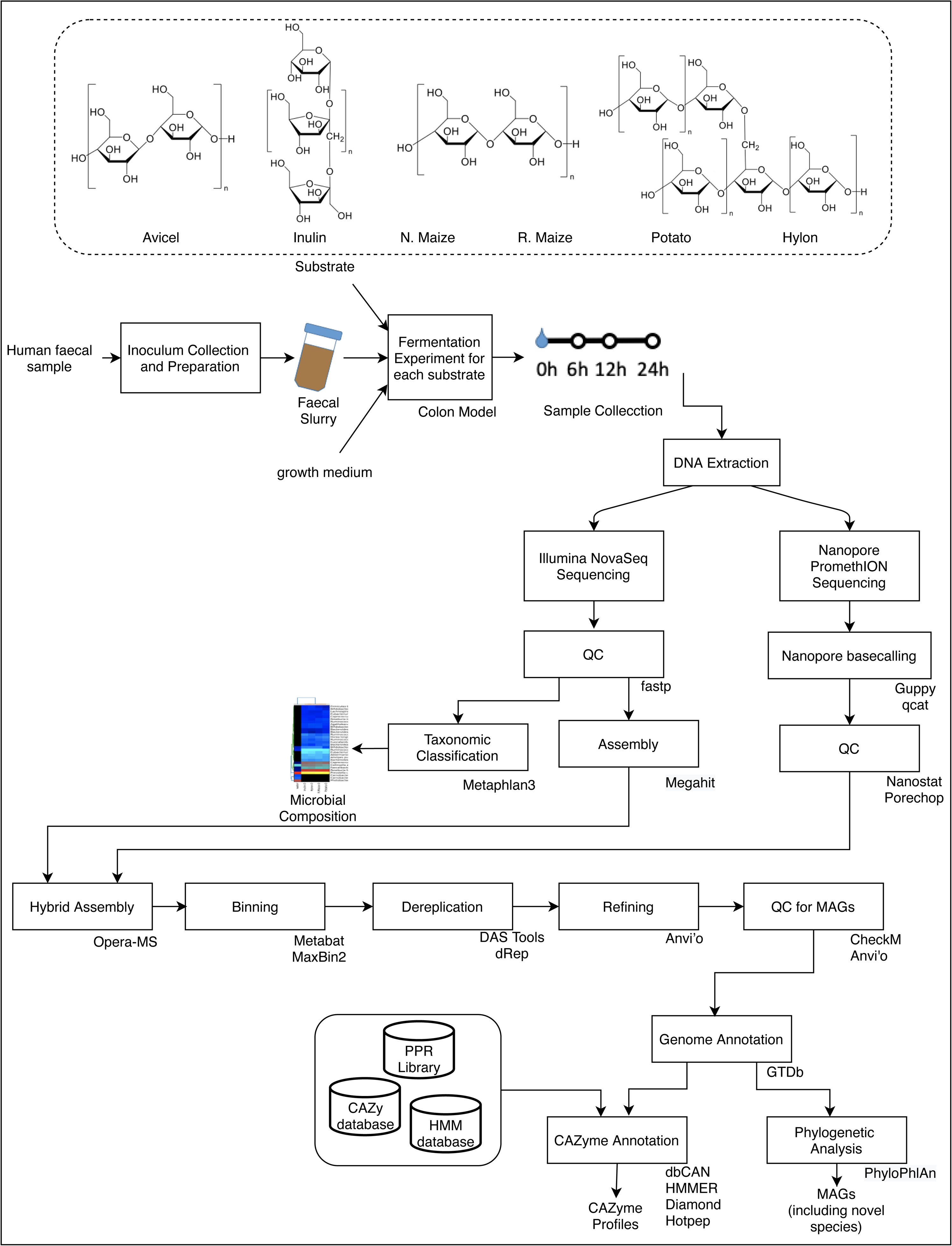
Workflow for bioinformatics analysis of combined Illumina NovoSeq and Oxford Nanopore PromethION metagenomics data collected in a model colon study of the fermentation of different carbohydrate substrates with contrasting structures (Avicel, Inulin, Normal maize (N.maize), Retrograded maize (R.maize), Potato and Hylon) by the gut microbiota present in a human stool sample.

**Figure 2.**
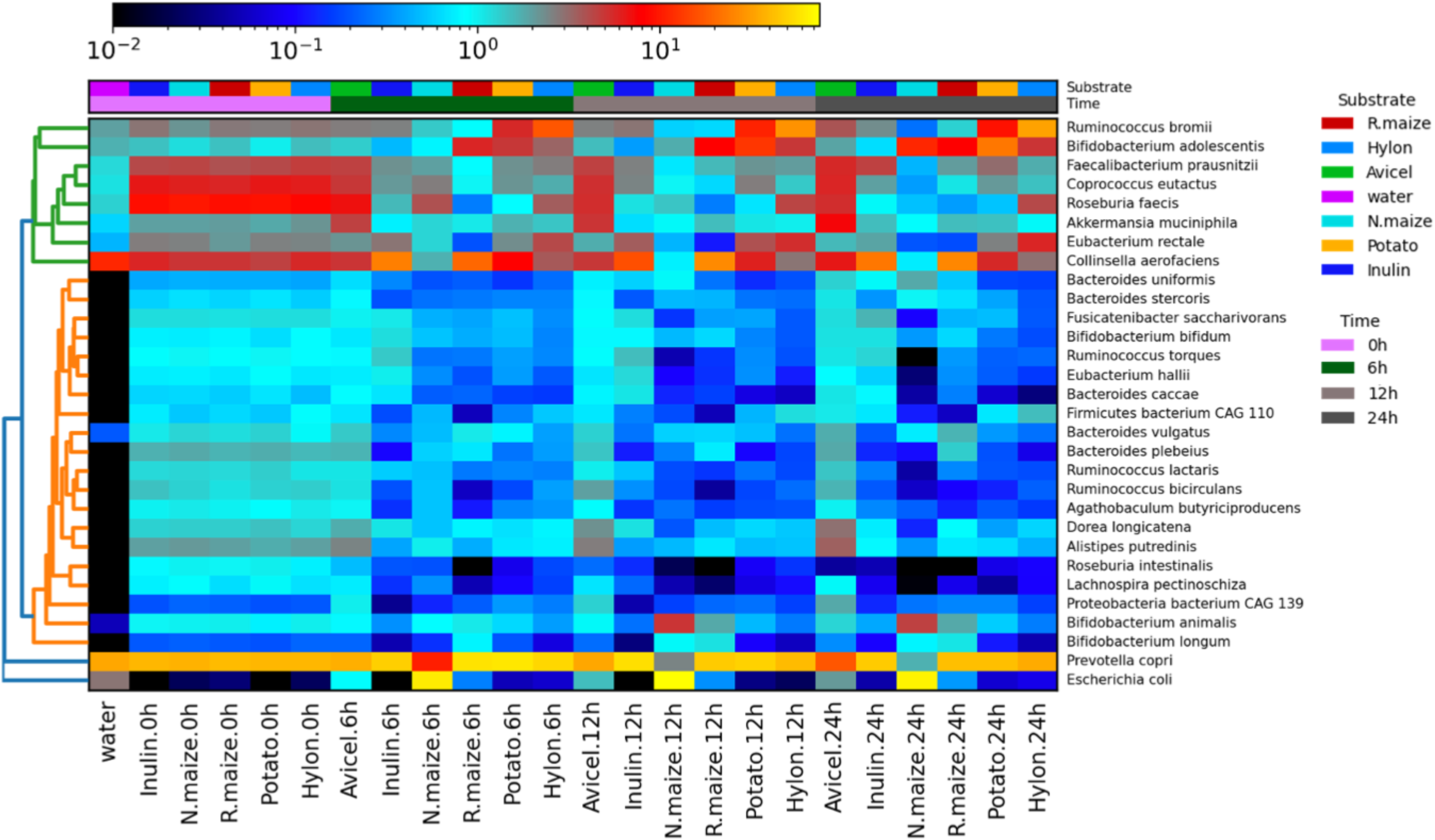
**Hierarchical clustering** of the top 30 selected gut microbial species present after fermentation of Avicel, Inulin, N.maize, R.maize, Potato and Hylon at 0h, 6h, 12h and 24h in the model colon. The hierarchical clustering also includes a water sample (“the kitome”).

#### PromethION sequencing

The two sequencing runs generated 144 giga base pairs (Gbp) of raw sequences. In the first run, all 23 samples were analysed while in the second batch, 12 samples from hylon, inulin and r.maize were selected. The first run produced 7.87 million reads with an average read length of 3419 ± 57 bp and the second run generated 21.6 million reads with an average read length of 4707 ± 206 bp. Consolidating the runs, trimming and quality filtering resulted in the removal of 33.3 ± 14.7 % of reads (Supplementary Table 1). Median read lengths after trimming were 4972.5 ± 229 bp and the median quality score was 9.7 ± 0.9.

#### Illumina sequencing

All 23 samples provided high quality sequences (Q value > 30) generating a mean of 27 million reads per sample. Quality and read length (<60 bp) filtering removed 2.96 % of reads (Supplementary Table 1).

### Dynamic shifts in taxonomic profiles among carbohydrate treatments

Hierarchical clustering for the taxonomic profiling using MetaPhlAn3 for each sample is shown in Supplementary Table 2. At baseline (0h), profiles of the top 30 selected species by clustering (using Bray-Curtis distances for samples and species, and a complete linkage) is similar for all treatments, as expected (**Error! Reference source not found.**). This uniform profile was distinct from the water control sample (a.k.a. ‘the kitome’). The water blank also had less than 3% (NovoSeq) and less than 0.2% (PromethION) of the reads of the samples. Microbiome shifts were apparent from 6h in the n.maize treatment which showed a very high abundance of *E. coli*, indicating contamination. After 12h, the profiles changed further with a higher abundance of *E. coli* and *B. animalis* in the n.maize treatment while the r.maize and inulin treatment profiles were similar, as were the potato and Hylon treatment profiles. By the last sampling point (24h), potato and hylon had similar profiles which are also similar to r.maize. The most abundant species in all the substrates was consistently Prevotella copri which decreased in abundance over time but remained one of the most abundant species throughout. After 6h and 12h, *Ruminococcus bromii* (a keystone starch degrader) and *Bifidobacterium adolescentis* increased in abundance in the r.maize, potato and Hylon treatments. *Faecalibacterium praunitzii* decreased in abundance in inulin at 6h and 12h and then increased in abundance for inulin and avicel at 24h.

Dynamic shifts in the microbiome were estimated using PCoA (Supplementary Figure 1), with 77% of total variance being explained by the first two components. As expected, the 0h profiles clustered closely together. The most distinct taxonomic change in microbial community composition was apparent in the Avicel treatment after 24h. Inulin and r.maize profiles clustered more closely together than potato and Hylon profiles. Inverse Simpson index results followed a similar pattern for changes in diversity, which decreased after 0h followed by a gradual increase (Supplementary Figure 2). However, in the Avicel treatment there was a different pattern of taxonomic shifts with a large number of taxa increasing in abundance after 12h.

### Hybrid metagenome assemblies vs short-read only assemblies

Using Opera-MS, we combined PromethION reads with Illumina assemblies to produce hybrid assemblies. The assembly statistics for short-read-only and hybrid assemblies are shown in Supplementary Table 3 and **Error! Reference source not found.**. The longest N50 and the largest contig per treatment were generated using hybrid assemblies as expected (figure 3b & 3c). The overall length of assembled sequences was similar for both approaches (Figure 3d).

**Figure 3:**
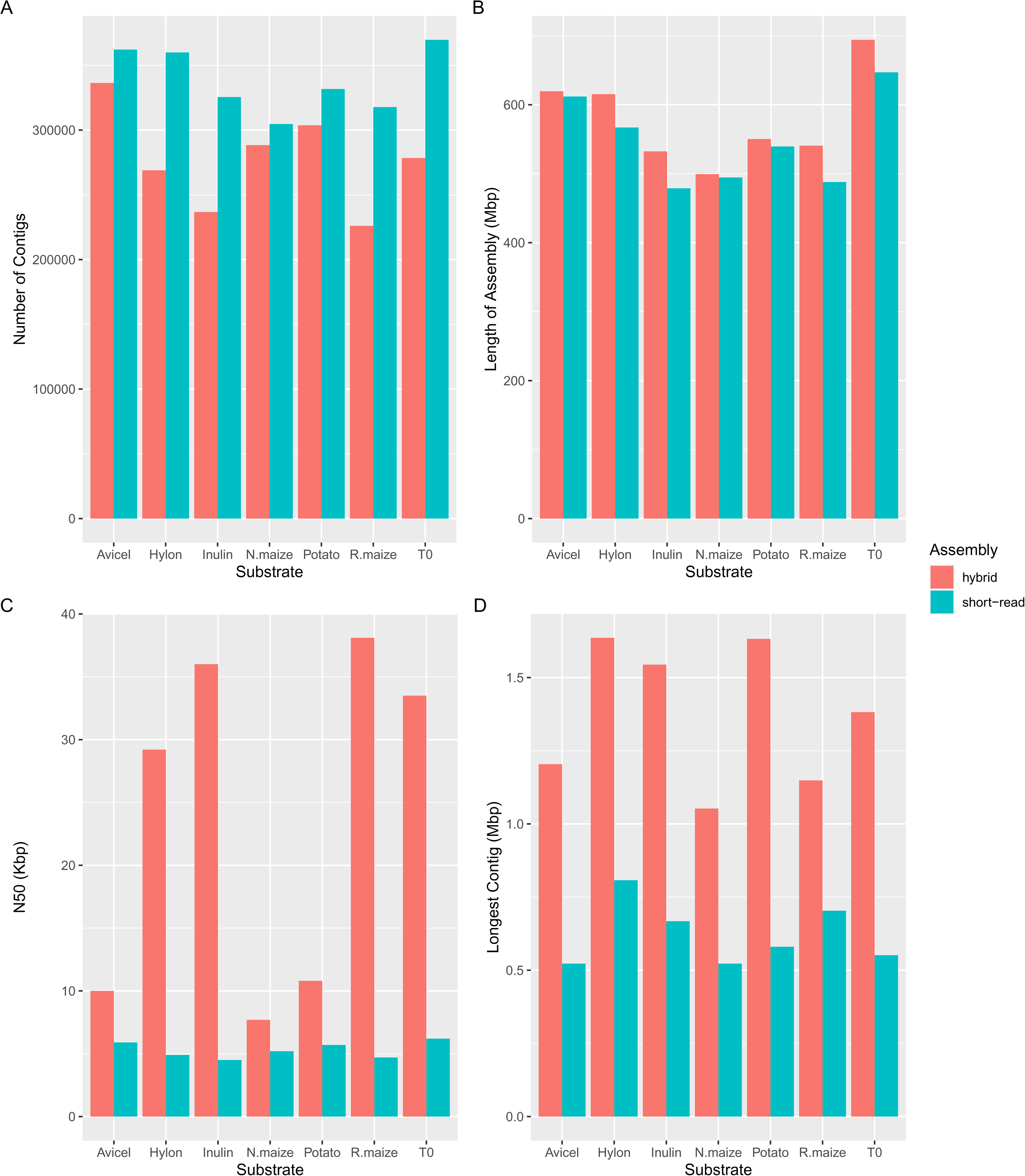
Comparison of Illumina short read assemblies and hybrid assemblies: a) shows the number of contigs per treatment, b) shows the N50, c) statistics on the largest contig, d) size of the total assembly for each carbohydrate treatment.

The reads from each treatment and collective T0 were co-assembled into hybrid assemblies and binned into MAGs. In total we binned and refined 509 MAGs that met the MIMAG quality score criteria[20] of which 65% (n=333) were high-quality (Figure 4; Supplementary table 4). From the co-assemblies, thirty-five MAGs had an N50 of > 500,000 Mbp and 158 MAGs were assembled into < 30 scaffolds. The MAGs were dereplicated into primary and secondary clusters according to Average Nucleotide identity (ANI) (primary clusters <97%; secondar clusters <99%). In total, we identified 151 MAG secondary clusters (Supplementary table 5). Each genome cluster consisted of between one and seven genomes based on their genome similarity.

**Figure 4:**
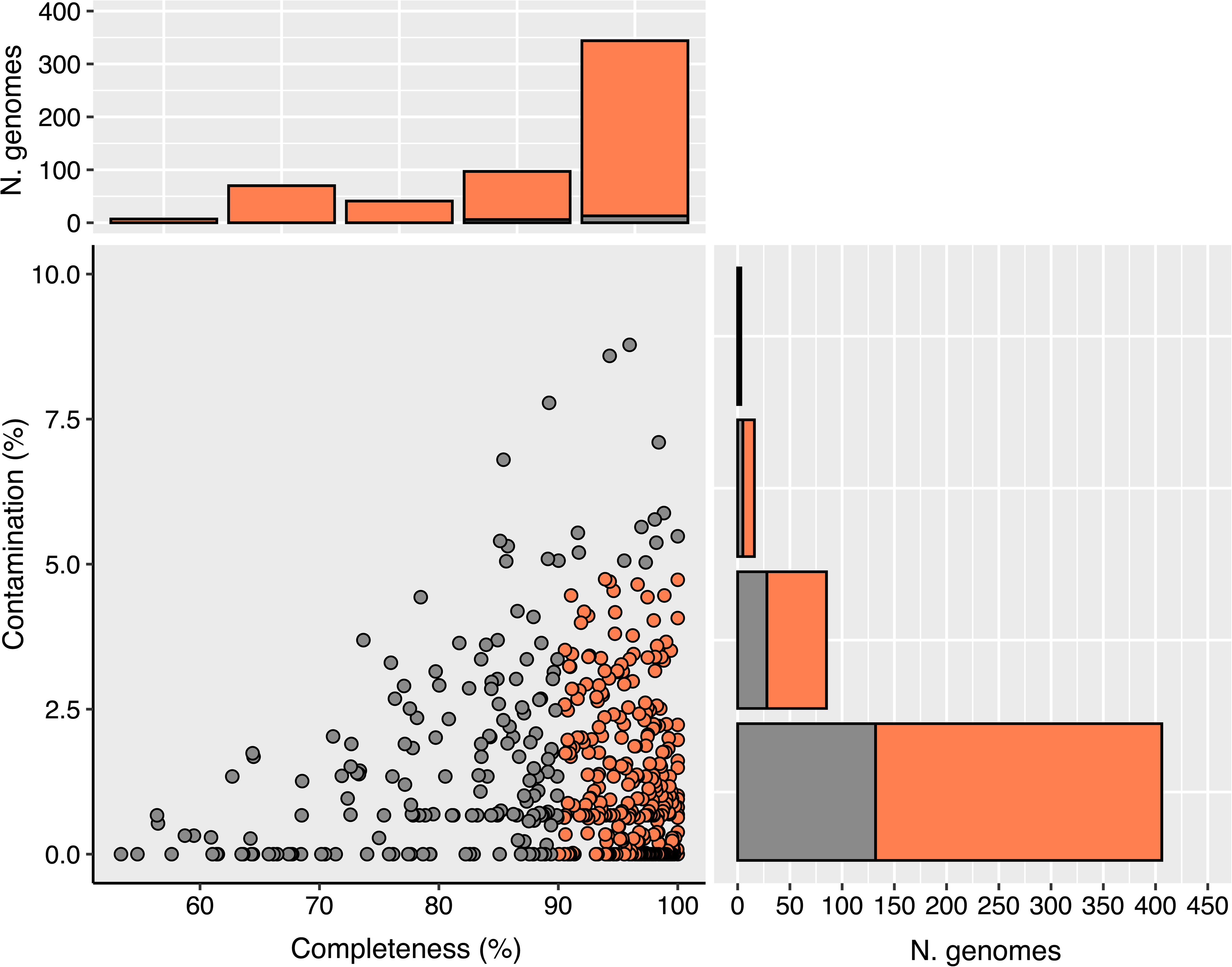
MAG quality. Dots represent each MAG. Completeness and contamination scores were estimated using CheckM. Colours are based on the MAG standards (high quality as >90% completeness & <5% contamination; good quality as <90%-60% completeness and >5% - 10% contamination. The horizontal and vertical bar charts provide the number of genomes with high completeness and low contamination scores.

### Taxonomic annotation of MAGs

Proposed bacterial taxonomy using GTDb was represented in existing bacterial families: All MAG clusters had > 99% identity to existing genera (Supplementary Table 6). Here, 49 of the 151 MAG clusters was named using alphanumeric genus names and 19 MAGs clusters was named using alphanumeric species names. In order to provide a clear and stable genus and species names, the MAGs were directly compared in NCBI to check whether these MAGs were already named or had any culture representatives. We identified 12 MAGs that was previously named and/or cultured so the Latin binomial for these MAGs was updated (Supplementary table 7). For the rest of the MAGs, we used the approach described in Pallen *et al*[21] to provide novel Latin names to 56 MAG clusters. We have provided 12 new genus names and 51 novel species names (Table 1). In addition, MAG assembly statistics for the MAGs in the present study was compared to the representative assemblies in GTDb (Supplementary Table 8). We found that while the average overall assembly length was almost similar (an average of 2,250,870 bp in the present study vs. 2,541,312bp in GTDb), there were far fewer contigs in our assemblies (an average of 67 contigs in the present study vs. 160 in GTDb), and therefore our MAGs may be considered to be of higher quality.

### Carbohydrate structure drives progression of bacterial diversity

Relative abundance of each MAG within treatments was calculated and log fold change of abundance between treatments was used to estimate change in relative abundance (Supplementary table 9). In total, 36 of 151 clusters exhibited ≥2-log fold increase in relative abundance for all treatments. Specifically, ≥2-log fold change in abundance was seen in 6, 12, 11 and 18 MAGs for Avicel, Hylon, potato and r.maize treatments, respectively (Figure 5). The genomes were partitioned as early (0h up to 6h) and late degraders (12h to 24h) according to when they first showed an increase in relative abundance (Supplementary Table 10).

**Figure 5:**
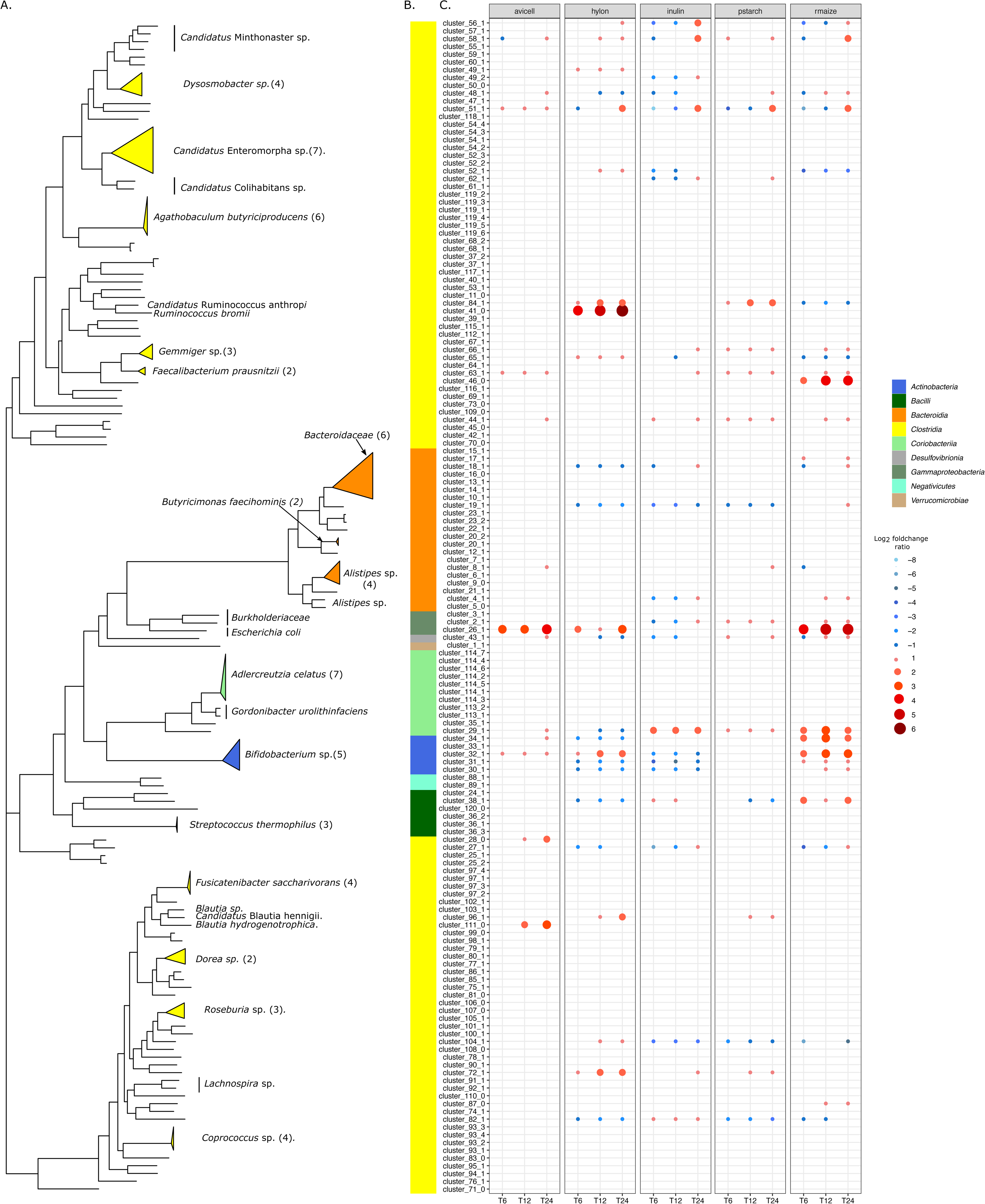
Phylogenomic tree and fold changes. The phylogenetic tree was **constructed from concatenated protein sequences using PhyloPhlAn and illustrated using ggtree.** Clades belonging to similar bacterial family and bacterial genus were collapsed. The colour strips represent the phylum-level distribution of the phylogenetic tree. Dot plot shows the decrease (negative log_2_ fold change; blue shades) and increase (positive log_2_ fold change; red shades) of read proportions from 0h to 6h, 0h to 12h and 0h to 24h for all treatments.

Relative abundance of all MAGs from each treatment was aggregated and plotted for each time period (Supplementary figure 3). Relative abundance was constant for Avicel throughout indicating low activity of the MAGs in utilising crystalline cellulose, likely reflecting the very limited fermentability of microcrystalline cellulose. As for other maize starches (hylon, r.maize and potato), the read proportions showed an overall reduction in abundance, with only starch degrading MAGs increasing in abundance.

### CAZyme family interplay with the carbohydrate treatments

For identifying CAZymes in the MAGs, genome-predicted proteins identified by Prodigal were compared with the CAZy database using dbCAN2 (Supplementary table 11). CAZyme counts specifically for Glycoside hydrolases (GH) and Carbohydrate binding modules (CBM) for all clusters showed a high representation of the profiles with GH13, GH2 and GH3 accounting for 34.1% of all counts (Supplementary Figure 4). CAZyme profiles for MAGs with > 2-log fold change are highlighted in Supplementary table 12 and Figure 6. Although six genomes were identified as associated with the degradation of cellulose, none contained any characteristic cellulose active CAZy proteins indicating multiple cross feeders. *Collinsella aerofaciens*_J (cluster 29_1), *Candidatus Minthovivens enterohominis* (cluster 81_1) are novel genomes that showed a 2x log -fold increase when in the presence of inulin and also harboured multiple copies of inulinases (GH32). *Bacteroides uniformis*, a known inulin degrader also contained multiple copies of GH32. We identified a large representation of the amylolytic (starch degrading) gene family GH13 in Hylon (counts= 88), potato (counts=50) and r.maize (counts=77) treatments. As expected, GH13 was weakly represented in Avicel (counts=19) and inulin (counts=29) treatments (Figure 6). The presence of GH13 in MAGs was closely associated with CBM48, which is commonly appended to starch degrading GH13 enzymes.[22] In total, we identified several novel degraders and previously discovered degraders of the different carbohydrate treatments which are highlighted in Supplementary table 10.

**Figure 6:**
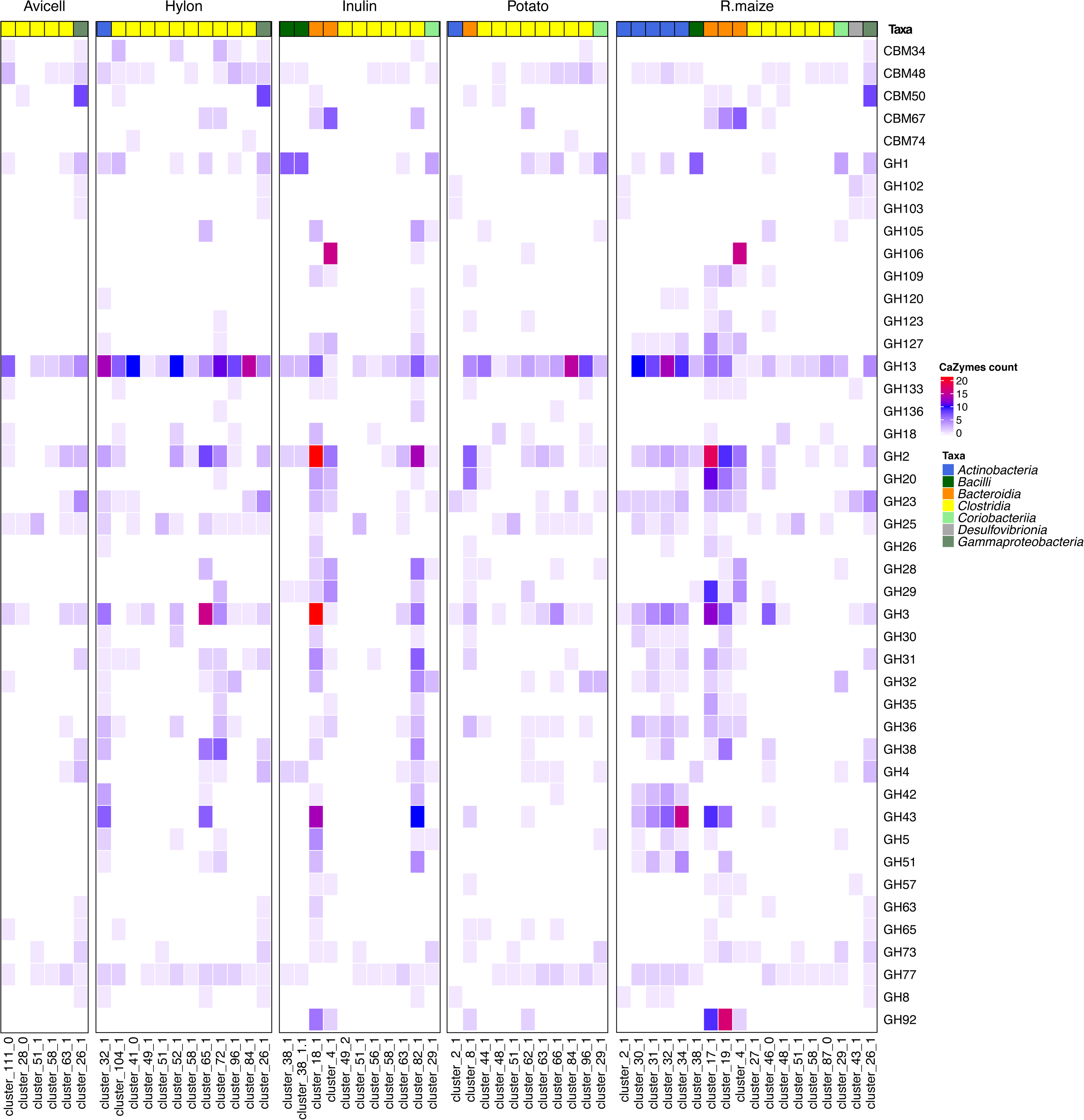
CAZyme profiles of selected-MAGs. The colour strip represents the phylum-based taxonomy annotation. The heat map represents the number of proteins identified for each CAZy protein family.

## Discussion

Using a hybrid assembly approach (i.e., combining NovaSeq short-read and PromethION long-read metagenomic data), we report species-level resolved taxonomic data identifying distinct changes in microbiome composition in response to different substrates. The large number of high-quality near-complete MAGs that we generated using this approach enabled us to functionally annotate the CAZymes in the MAGs and identify potential carbohydrate degrading species. Several of these species have not previously been identified as playing a role in starch fermentation (Figure 6 and Supplementary Table 9).

### High quality DNA for long-read sequencing was extracted using a bead beating protocol

The N50 for the PromethION reads was 4,972 bp, which is comparable with another recent study using bead-beating-based DNA extraction and provided adequate read lengths to be useful for assembly of MAGs.[17] A recent publication by Moss *et al.*[23] and associated protocol paper[24] suggested that bead beating DNA extraction protocols were unsuitable for long-read sequencing as they led to excessive shearing of DNA and therefore enzymatic cell lysis followed by phenol-chloroform purification were preferred to recover high molecular weight (HMW) DNA. This was not reflected in our experience. The N50’s obtained by Moss *et al.* for sequencing DNA extracted from stool samples by phenol-chloroform on the PromethION platform ranged from 1,432 bp to 5,205 bp, which on average was shorter than the N50 we obtained using comparable samples extracted by a bead beating protocol. This is in agreement with Bertrand et al.[14] who directly compared commercial bead beating and phenol-chloroform extraction protocols for extracting HMW DNA from stool samples for MinION sequencing and found that while phenol-chloroform gave higher molecular weights of DNA, the DNA was of low integrity compromising sequencing quality.

### Hybrid assemblies allow generation of near complete MAGs

We found larger N50s and longest contigs when using hybrid assemblies compared with short-read assemblies; this is in agreement with previous benchmarking data using a combined MinION and Illumina hybrid approach to sequence mock communities, human gut samples,[14] and rumen gut microbiota samples.[17] This allowed us to assemble 509 MAGs across all the major phylogenetic groups (Supplementary file 5), with representatives from ten bacterial classes and 28 families, including both Gram-positive and Gram-negative species. Bertand et al.[14] found that phenol-chloroform extractions led to underrepresentation of ‘hard to sequence’ gram-positive species such as those of the genus *Bifidobacterium*. In the present study near-complete MAG’s were recovered from 5 different species of *Bifidobacterium*, in contrast to Moss *et al.*[23] who were unable to recover *Bifidobacterium* MAG’s from the PromethION data produced using their enzyme and phenol-chloroform based extraction method (although they were able to recover Bifidobacterium MAG’s from short-read data which was obtained following a bead beating based DNA extraction of the same samples). This indicates that bead beating is necessary to obtain accurate representations of the microbial community in human stool samples. The bead beating DNA extraction protocol used in this study was also recommended by the Human Microbiome Project to avoid biases in microbiome samples.[25, 26]

We have provided *Candidatus* names to 70 bacterial species which do not currently have representative Latin binomial names in the GTDB database (Table 1 and Supplementary Table 7). Our decision to provide names for these species reflects the higher quality of MAGs compared to those currently represented in the databases (Supplementary Table 8).

### Structural diversity in substrates drives changes in microbial communities

Over the 24h fermentation, microbial communities rapidly diverged depending on substrate. The smallest change in community composition occurred in the Avicel treatment, as would be expected, given that Avicel was the most recalcitrant substrate evaluated, with very limited fermentability.[27] Each substrate resulted in distinct changes in microbial community composition, supporting previous findings that chemically-identical but structurally-diverse starches can result in distinct changes in microbial community composition.[7, 8]

### Changes in microbial composition are related to the ability to degrade structurally diverse substrates

To better understand potential mechanisms driving the changes in microbial species composition in response to different substrates, we explored the CAZyme profiles of our microbial community.[28] We found the greatest number and diversity of CAZyme genes were in the genomes of *Bacteroidetes* (Figure 6 and Supplementary Figure 4), as has previously been computationally estimated for the human gut microbiome.[10, 29] This is in contrast to rumen microbiomes where *Fibrobactares* are the primary fibre-degrading bacterial group.[17]

We identified genomes that increased in abundance during either early or late stages of fermentation suggesting that their involvement in substrate degradation was either as primary (early) or secondary (late) degraders (Figure 5). We also identified differences in abundance of particular CAZyme-encoding genes amongst species which may reflect their specialisation to specific substrates (Figure 6). *Bacteroides uniformis* has been characterised as an inulin-degrading species,[30] and in our analysis it was identified during inulin fermentation and had three copies of the GH32 (inulinase) gene and a gene encoding the inulin binding domain, CBM38. *Candidatus Minthovivens enterohominis* also increased in abundance early in inulin degradation, and its genome contained five copies of the GH32 gene. *Faecalibacterium prausnitzii* increased in abundance with inulin supplementation and has been shown to have the ability to degrade inulin when co-cultured with primary degrading species.[31, 32] *F. prausnitzii* was also found to increase in abundance for cellulose, but not for the starch based substrates.

Avicel is a highly crystalline cellulose that is resistant to fermentation; the human gut microbiota has a very limited capacity to degrade celluloses.[33] Interestingly, the largest increase in abundance we observed was for *Blautia hydrogenotrophica*; which has been reported in association with cellulose fermentation since it acts as an acetogen using hydrogen produced by primary degraders of cellulose.[34]

In all starch treatments, there were large increases in the proportion of identified genes that encoded GH13 (the major amylolytic gene family including α-amylase, α-glucosidase and pullulanase) reflecting selection for starch-degrading species (Figure 6); this was also the case for CBM48 which is also involved in starch degradation (Figure 6).[22] Our analysis identified several well-known starch degrading species, most notably *R. bromii and B. adoloscentis* (Figure 5). *R. bromii* is a well characterised specialist on highly recalcitrant starch,[35] possessing specialised starch-degrading machinery termed the ‘amylosome’; it was only identified in the most recalcitrant starch treatments (Hylon and potato). Previous genome sequencing of an *R. bromii* isolate reported 15 GH13 genes;[35] 14 GH13 genes were identified in the *R. bromii* MAG assembled in this study. In the potato treatment another closely related but less well characterised *Rumminococcus* species with ten GH13 genes and one CBM48 gene was identified.

A previously uncultured *Blautia* species was identified possessing eight GH13 and three CBM48 genes which increased in abundance in response to Hylon and potato. *Blautia* species have previously been shown to increase in abundance in response to resistant starch.[36, 37] We also identified four further previously-uncharacterised species that increased in abundance and had more than five GH13 genes: *Candidatus Cholicenecus caccae, Candidatus Eisenbergiella faecalis, Candidatus Enteromorpha quadrami and Candidatus Aphodonaster merdae*.

Maize starch treatments (r.maize and Hylon) showed increases in abundance of *Bifidobacterium* species. Previous studies have characterized *Bifidobacterium* as a starch-degrading genus.[38] The only *Bifidobacterium* species to increase in abundance in response to Hylon was *B. adolescentis*, which is known to utilise to this hard-to-digest starch better than other *Bifidobacterium* species,[39]; a broader range of *Bifidobacterium* species increased in abundance in response to the more accessible r.maize.

## Conclusion

We have demonstrated that deep long- and short-read metagenomic sequencing and hybrid assembly has great potential for studying the human gut microbiota. We identified species-level resolved changes in microbial community composition and diversity in response to carbohydrates with different structures over time, identifying succession of species within the fermenter. To provide functional information about these species we obtained over 500 MAGs from a single human stool sample. Annotating CAZyme genes in MAGs from species enriched for by fermentation of different carbohydrates allowed us to identify species specialised in degradation of defined carbohydrates, increasing our knowledge of the range of species potentially involved in starch metabolism in the human gut.

## Material and Methods

A schematic overview of the workflow and experimental design is displayed in Figure 1. **Substrates**. Native maize starch (catalogue no. S4126), native potato starch (catalogue no. 2004), Avicel PH-101 (catalogue no. 11365) and chicory inulin (catalogue no. I2255) were purchased from Sigma-Aldrich, (Gillingham, UK). Hylon VII® was kindly provided as a gift by Ingredion Incorporated (Manchester, UK).

Retrograded maize starch was prepared from 40g of native maize starch in 400 mL of deionized water. The slurry was stirred continuously at 95°C in a water bath for 20 minutes. The resulting gel was cooled to room temperature for 60 minutes, transferred to aluminium pots (150 mL, Ampulla, Hyde UK), and stored at 4°C for 48 hours. The retrograded gel was then frozen at −80°C for 12 hours and freeze-dried (LyoDry, MechaTech Systems Ltd, Bristol, UK) for 72 hours.

Each substrate (0.500 ± 0.005g, dry weight) was weighed in sterilized fermentation bottles (100 mL) prior to start of the experiment.

### Inoculum collection and preparation

A single human faecal sample was obtained from one adult (≥ 18 years old), free-living, healthy donor who had not taken antibiotics in the 3 months prior to donation and was free from gastrointestinal disease. Ethical approval was granted by Human Research Governance Committee at the Quadram Institute (IFR01/2015) and London - Westminster Research Ethics Committee (15/LO/2169) and the trial was registered on clinicaltrials.gov (NCT02653001). A signed informed consent was obtained from the participant prior to donation. The stool sample was collected by the participant, stored in a closed container under ambient conditions, transferred to the laboratory and prepared for inoculation within 2 hours of excretion. The faecal sample was diluted 1:10 with pre-warmed, anaerobic, sterile phosphate buffer saline (0.1M, pH 7.4) in a double meshed stomacher bag (500 mL, Seward, Worthing, UK) and homogenized using a Stomacher 400 (Seward, Worthing, UK) at 200 rpm for two cycles, each of 60 seconds length.

### Batch fermentation in the model colon

Fermentation vessels were established with media adapted from Williams *et al.*,[40] In brief, each vessel (100 mL) contained an aliquot (3.0 mL) of filtered faecal slurry, 82 mL of sterilized growth medium, and one of the six substrates for experimental evaluation: native Hylon VII or native potato starch (highly recalcitrant starches); native maize starch or gelatinized, retrograded maize starch (accessible starches); Avicel PH-101 (insoluble fibre; negative control); and chicory inulin (fermentable soluble fibre; positive control). There was also a media only control with no inoculum (blank) making a total of seven fermentation vessels.

For each fermentation vessel the growth medium contained 76 mL of basal solution, 5 mL vitamin phosphate and sodium carbonate solution, and 1 mL reducing agent. The composition of the various solutions used in the preparation of the growth medium is described in detail in **Supplementary Table 13**. A single stock (7 litres) of growth medium was prepared for use in all vessels. Vessel fermentations were pH controlled and maintained at pH 6.8 to 7.2 using 1N NaOH and 1N HCl regulated by a Fermac 260 (Electrolab Biotech, Tewkesbury, UK). A circulating water jacket maintained the vessel temperature at 37°C. Magnetic stirring was used to keep the mixture homogenous and the vessels were continuously sparged with nitrogen (99% purity) to maintain anaerobic conditions. Samples were collected from each vessel at 0 (5 min), 6, 12, and 24 hours after inoculation. The biomass from two 1.8 mL aliquots from each sample were concentrated by refrigerated centrifugation (4°C; 10,000 g for 10 min), the supernatant removed, and the pellets stored at −80°C prior to bacterial enumeration and DNA extraction; one pellet was used for enumeration and one for DNA extraction.

### Bacterial cell enumeration

All materials used for bacterial cell enumeration were purchased from Sigma-Aldrich (Gillingham, UK), unless specified otherwise. To each frozen pellet, 400 µL of PBS and 1100 µL of 4% paraformaldehyde (PFA) were added and gently thawed at 20°C for 10 minutes with gentle mixing. Once thawed, each resuspension was thoroughly mixed and incubated overnight at 4°C for fixation to occur. The resuspensions were then centrifuged for 10 minutes at 8000 x g, the supernatant removed, and the residual pellet washed with 1 mL 0.1% Tween-20. This pellet then underwent two further washes in PBS to remove any residual PFA and was then resuspended in 600 µL PBS: ethanol (1:1).

The fixed resuspensions were centrifuged for 3 minutes at 16000 x g, the supernatant removed, and the pellet resuspended in 500 µL 1 mg/mL lysozyme (100 µL 1M Tris HCl at pH 8, 100 µL 0.5 M EDTA at pH 8, 800 μL water, and 1 mg lysozyme, catalogue no. L6876) and incubated at room temperature for 10 minutes. After thorough mixing and centrifugation for 3 minutes at 16000 x g, the supernatant was removed, and the pellet washed with PBS. The resulting pellet was then resuspended in 150 µL of hybridisation buffer (HB, per mL: 180 µL 5 M NaCl, 20 µL 1M Tris HCl at pH 8, 300 µL Formamide, 499 µL water, 1 µL 10% SDS), centrifuged, the supernatant removed and the remaining pellet resuspended again in 1500 µL of HB and stored at 4°C prior to enumeration. For bacterial enumeration, 1 µL of Invitrogen SYTO 9 (catalogue no. S34854, Thermo Fisher Scientific, Loughborough, UK) was added to 1 mL of each fixed and washed resuspension. Within 96-well plate resuspensions were diluted to 1:1000 and the bacterial populations within them enumerated using flow cytometry (Luminex Guava easyCyte 5) at wavelength of 488nm and Guava suite software, version 3.3.

### DNA extraction

Each pellet was resuspended in 500 μL (samples collected at 0 and 6 hr) or 650 μL (samples collected at 12 and 24 hr) with chilled (4°C) nuclease-free water (Sigma-Aldrich, Gillingham, UK). The resuspensions were frozen overnight at −80°C, thawed on ice and an aliquot (400 μL) used for bacterial genomic DNA extraction. FastDNA® Spin Kit for Soil (MP Biomedical, Solon, US) was used according to the manufacturer’s instructions which included two bead-beating steps of 60s at a speed of 6.0m/s (FastPrep24, MP Biomedical, Solon, USA). DNA concentration was determined using the Quant-iT™ dsDNA Assay Kit, high sensitivity kit (Invitrogen, Loughborough, UK) and quantified using a FLUOstar Optima plate reader (BMG Labtech, Aylesbury, UK).

### Illumina NovaSeq Library preparation and sequencing

Genomic DNA was normalised to 5 ng/µL with elution buffer (10mM Tris-HCl). A miniaturised reaction was set up using the Nextera DNA Flex Library Prep Kit (Illumina, Cambridge, UK). 0.5 µL Tagmentation Buffer 1 (TB1) was mixed with 0.5 µL Bead-Linked Transposomes (BLT) and 4.0 µL PCR-grade water in a master mix and 5 µL added to each well of a chilled 96-well plate. 2 µL of normalised DNA (10 ng total) was pipette-mixed with each well of tagmentation master mix and the plate heated to 55°C for 15 minutes in a PCR block. A PCR master mix was made up using 4 µL kapa2G buffer, 0.4 µL dNTP’s, 0.08 µL Polymerase and 4.52 µL PCR grade water, from the Kap2G Robust PCR kit (Sigma-Aldrich, Gillingham, UK) and 9 µL added to each well in a 96-well plate. 2 µL each of P7 and P5 of Nextera XT Index Kit v2 index primers (catalogue No. FC-131-2001 to 2004; Illumina, Cambridge, UK) were also added to each well. Finally, the 7 µL of Tagmentation mix was added and mixed. The PCR was run at 72°C for 3 minutes, 95°C for 1 minute, 14 cycles of 95°C for 10s, 55°C for 20s and 72°C for 3 minutes. Following the PCR reaction, the libraries from each sample were quantified using the methods described earlier and the high sensitivity Quant-iT dsDNA Assay Kit. Libraries were pooled following quantification in equal quantities. The final pool was double-SPRI size selected between 0.5 and 0.7X bead volumes using KAPA Pure Beads (Roche, Wilmington, US). The final pool was quantified on a Qubit 3.0 instrument and run on a D5000 ScreenTape (Agilent, Waldbronn, DE) using the Agilent Tapestation 4200 to calculate the final library pool molarity. qPCR was done on an Applied Biosystems StepOne Plus machine. Samples quantified were diluted 1 in 10,000. A PCR master mix was prepared using 10 µL KAPA SYBR FAST qPCR Master Mix (2X) (Sigma-Aldrich, Gillingham, UK), 0.4 µL ROX High, 0.4 µL 10 μM forward primer, 0.4 µL 10 μM reverse primer, 4 µL template DNA, 4.8 µL PCR grade water. The PCR programme was: 95°C for 3 minutes, 40 cycles of 95°C for 10s, 60°C for 30s. Standards were made from a 10 nM stock of Phix, diluted in PCR-grade water. The standard range was 20 pmol, 2 pmol, 0.2 pmol, 0.02 pmol, 0.002 pmol, 0.0002 pmol. Samples were then sent to Novogene (Cambridge, UK) for sequencing using an Illumina NovaSeq instrument, with sample names and index combinations used. Demultiplexed FASTQ’s were returned on a hard drive.

### Nanopore library preparation and PromethION sequencing

Library preparation was performed using SQK-LSK109 (Oxford Nanopore Technologies, Oxford, UK) with barcoding kits EXP-NBD104 and EXP-NBD114. The native barcoding genomic DNA protocol by Oxford Nanopore Technologies (ONT) was followed with slight modifications. Starting material for the End-Prep/FFPE reaction was 1 μg per sample in 48 μL volume. 3.5 µL NEBNext FFPE DNA Repair Buffer (NEB, New England Biolabs, Ipswich, USA), 3.5 µL NEB Ultra II End-prep Buffer, 3 µL NEB Ultra II End-prep Enzyme Mix and 2 µL NEBNext FFPE DNA Repair Mix (NEB) were added to the DNA (final volume 60 µL), mixed slowly by pipetting and incubated at 20°C for 5 minutes and then 65°C for 5 minutes. After a 1X bead wash with AMPure XP beads (Agencourt, Beckman Coulter, High Wycombe, UK), the DNA was eluted in 26 µL of nuclease-free water. 22.5 µL of this was taken forward for native barcoding with the addition of 2.5 µL barcode and 25 µL Blunt/TA Ligase Master Mix (NEB) (final volume 50 µL). This was mixed by pipetting and incubated at room temperature for 10 minutes. After another 1X bead wash (as above), samples were quantified using Qubit dsDNA BR Assay Kit (Invitrogen, Loughborough, UK). In the first run, samples were equimolar pooled to a total of 900 ng in a volume of 65 µL. In the second run, samples were pooled to 1700 ng followed by a 0.4X bead wash to achieve the final volume of 65 µL. 5 µL Adapter Mix II (ONT), 20 µL NEBNext Quick Ligation Reaction Buffer (5X) and 10 µL Quick T4 DNA Ligase (NEB) were added (final volume 100 µL), mixed by flicking, and incubated at room temperature for 10 minutes. After bead washing with 50 µL of AMPure XP beads and two washes in 250 µL of Long Fragment Buffer (ONT), the library was eluted in 25 µL of Elution Buffer and quantified with Qubit dsDNA BR and TapeStation 2200 using a Genomic DNA ScreenTape (Agilent Technologies, Edinburgh, UK). 470 ng of DNA was loaded for sequencing in the first run and 400 ng in the second run. The final loading mix was 75 µL SQB, 51 µL LB and 24 µL DNA library.

Sequencing was performed on a PromethION Beta using FLO-PRO002 PromethION Flow Cells (R9 version). The sequencing runtime was 57 hours for Run 1 and 64 hours for Run 2. Flow cells were refuelled with 0.5X SQB (75 µL SQB and 75 µL nuclease free water) 40 hours into both runs.

### Bioinformatics analysis

The bioinformatics analysis was performed using default options unless specified otherwise.

#### Nanopore basecalling

Basecalling was performed using Guppy version 3.0.5+45c3543 (ONT) in high accuracy mode (model dna_r9.4.1_450bps_hac), and demultiplexed with qcat version 1.1.0 (Oxford Nanopore Technologies, https://github.com/nanoporetech/qcat). Sequence quality: For Nanopore, sequence metrics were estimated by Nanostat version 1.1.2[41]. In total, 22 million sequences were generated with a median read length of 4500 bp and median quality of 10 (phred). Quality trimming and adapter removal was performed using Porechop version 0.2.3 (https://github.com/rrwick/Porechop). For Illumina, quality control was done for paired-end reads using fastp, version 0.20.0.[42] to remove adapter sequences and filter out low-quality (phred quality < 30) and short reads (length < 60 bp). After quality control, the average number of reads in the samples was over 26.1 million reads, with a minimum of 9.7 million reads; the average read length was 148 bp. *Taxonomic profiling:* Trimmed and high-quality short reads are processed using MetaPhlAn3 version 3.0.2,[43] to estimate both microbial composition to species level and also the relative abundance of species from each metagenomic sample. MetaPhlAn3 uses the latest marker information dataset, CHOCOPhlAn 2019, which contains ∼1 million unique clade-specific marker genes identified from ∼100,000 reference genomes; this includes bacterial, archaeal and eukaryotic genomes. Hclust2 was used to plot the hierarchical clustering of the different taxonomic profiles at each time point [https://github.com/SegataLab/hclust2]. The results of the microbial taxonomy were analysed in RStudio Version 1.1.453 (http://www.rstudio.com/).

Principle Coordinate analyses using the pcoa function in the ape package version 5.3 (https://www.rdocumentation.org/packages/ape/versions/5.3) and the vegan package was used to identify differences in microbiome profiles amongst treatments.

#### Hybrid assembly

Trimmed and high-quality Illumina reads were merged per treatment, and then used in a short-read-only assembly using Megahit version 1.1.3.[44, 45] Then OPERA-MS[46] version 0.8.2, was used to combine the short-read only assembly with high-quality long reads, to create high-quality hybrid assemblies. By combining these two technologies, OPERA-MS overcomes the issue of low-contiguity of short-read-only assemblies and the low base-pair quality of long-read-only assemblies.

#### Genome binning, quality, dereplication and comparative genomics of hybrid assemblies

The hybrid co-assemblies from Opera-MS[46] were used for binning. Here, Illumina reads for each time period were mapped to the co-assembled contigs to obtain a coverage map. Bowtie2 version 2.3.4.1 was used for mapping, and samtools to convert SAM to BAM format. MaxBin2 version 2.2.6[47] and MetaBat2 version 2.12.1[48] which uses sequence composition and coverage information, was used to bin probable genomes using default parameters. The binned genomes and co-assembled contigs were integrated into Anvi’o version 6.1 for manual refinement and visual inspection of problematic genomes.[49] In particular, we used the scripts: ‘anvi-interactive’ to visualise the genome bins; ‘anvi-run-hmms’ to estimate genome completeness and contamination; ‘anvi-profile’ to estimate coverage and detection statistics for each sample; and ‘anvi-refine’ to manually refine the genomes. All scripts were run using default parameters. Additionally, DAS tool version 1.1.2 [50] was used to aggregate high-quality genomes from each treatment by using single copy gene-based scores and genome quality metrics to produce a list of good quality genomes for every treatment. Additionally, checkM version 1.0.18[51] was used on all final genomes to confirm completion and contamination scores. In general, genomes with a ‘quality satisfying completeness - 5*contamination > 50 score’ and/or with a ‘>60% completion and <10% contamination’ score according to CheckM, were selected for downstream analyses.

#### Dereplication into representative clusters

In order to produce a dereplicated set of genomes across all treatments, dRep version 2.5.0[52] was used. Pairwise genome comparisons or Average Nucleotide Identity (ANI) was used for clustering. dRep clusters genomes with ANIs of 97% were regarded as primary clusters, and genomes with ANI of 99 % regarded as secondary clusters. A representative genome is provided for each of the secondary clusters.

#### Relative abundance of genomes

Since co-assemblies were used for binning, relative abundance was calculated as the proportion of reads recruited to that bin across all time periods for each treatment. This provides an estimate of which time period recruited the most reads. To provide this estimate in relative terms, the value is normalised to the total number of reads that was recruited for that genome. As for Avicel that misses the time 0h, a mean relative abundance from each MAG in the cluster at time 0h was used. The relative abundance scores was provided by ‘anvi-summarize’ (from the Anvi’o package) as relative abundance. Further, fold changes were calculated between the relative abundance at time 0h to the corresponding relative abundance at 6h, 12h and 24h using gtools R package version 3.5.0. Fold changes provide an estimate of change in MAG abundance which might be a result from utilisation of a particular carbohydrate. Fold changes were converted to log ratios. MAGs with a fold change of 2x (log_2_ foldchange=1) were regarded as an active carbohydrate utiliser.

#### Metagenomic assignment and phylogenetic analyses

Genome bins that passed quality assessment were analysed for their closest taxonomic assignment. To assign taxonomic labels, the genome set was assigned into the microbial tree of life using GTDB version 0.3.5 and database R95 to identify the closest ancestor and obtain a putative taxonomy assignment for each genome bin. For genomes where the closest ancestor could not be determined, the Relative Evolutionary Distance (RED) to the closest ancestor and novel taxa names were provided. Using these genome bins, a phylogenetic tree was constructed using Phylophlan version 0.99 and visually inspected using iTOL version 4.3.1 and ggtree from package https://github.com/YuLab-SMU/ggtree.git. The R packages ggplot2 version 3.3.2, dplyr version 1.0.2, aplot, ggtree version 2.2.4 and inkscape version 1.0.1 were used for illustrations

#### Carbohydrate metabolism analyses

All representative genome clusters were annotated for CAZymes using dbCAN.[53] The genome’s nucleotide sequences were processed with Prodigal to predict protein sequences, and then three tools were used for automatic CAZyme annotation: a) HMMER[54] to search against the dbCAN HMM (Hidden Markov Model) database; b) DIAMOND[55] to search against the CAZy pre-annotated CAZyme sequence database; and c) Hotpep[56] to search against the conserved CAZyme PPR (peptide pattern recognition) short peptide library. To improve annotation accuracy, a filtering step was used to retain only hits to CAZy families found by at least two tools. The R packages ggplot2, dplyr, ComplexHeatmap version 2.4.3 and inkscape were used for illustrations.

## Acknowledgements

We thank Dave J. Baker for assisting with sequencing and the anonymous donor who provided faecal material for this study. We thank Dr. Judith Pell for assistance with editing the manuscript. We acknowledge the kind assistance of Prof. Aharon Oren for checking and correcting the grammar of the protologue species names.

## Author contributions

All authors read and contributed to the manuscript. AR, PR and JAJ are joint first authors. FJW conceived and designed the study. AR led on the preparation of the manuscript. AA and GLK prepared the sequencing libraries and did the sequencing. AR and PR did the sequence and bioinformatics analysis. TLV did the post-sequencing analysis. JAJ, KC and SH did the model colon experiments and DNA extractions. HH enumerated the bacterial cells. RG and MJP assisted with bioinformatic analysis and taxonomic descriptions. JOG provided long-read sequencing and molecular biology expertise; AJP provided bioinformatics expertise; and FJW provided expertise in carbohydrate structure and model colon protocols. FJW, JOG, AJP secured funding, provided management oversight and scientific direction.

## Ethical approval

Ethical approval was granted by the Human Research Governance Committee at the Quadram Institute (IFR01/2015) and the London - Westminster Research Ethics Committee (15/LO/2169). The trial is registered on clinicaltrials.gov (NCT02653001). A signed informed consent was obtained from the participant prior to donation.

## Funding statements

The authors gratefully acknowledge the support of the Biotechnology and Biological Sciences Research Council (BBSRC). This research was funded by: the BBSRC Institute Strategic Programme (ISP) Food Innovation and Health BB/R012512/1 and its constituent projects (BBS/E/F/000PR10343, BS/E/F/000PR10346); the BBSRC ISP Microbes in the Food Chain BB/R012504/1 and its constituent projects (BBS/E/F/000PR10348, BBS/E/F/000PR10349, BBS/E/F/000PR10352); and the BBSRC Core Capability Grant (BB/CCG1860/1). The funders had no role in study design, data collection and analysis, decision to publish, or preparation of the manuscript.

## Availability of data and materials

Raw read data from the PromethION and NovoSeq sequencing runs can be accessed through the NCBI SRA project number PRJNA722408 and can be accessed at https://dataview.ncbi.nlm.nih.gov/object/PRJNA722408?reviewer=ts65d8lkvj8nbv4mpfsar7sv3g. GenBank accession numbers for individual MAG’s within this ProjectID can be found in Supplementary Table 5.

## Competing interests

The authors declare that they have no competing interests

## Supplementary Files

**Supplementary Table 1:** Read stats and quality metrics for PromethION and Illumina sequence data

**Supplementary Table 2:** Taxonomy profiles of relative abundances for all treatments using MetaPhlAn3.

**Supplementary Table 3:** Assembly stats for short read assemblies using Megahit and hybrid assemblies using OPERA-MS

**Supplementary Table 4:** MAG genomic stats, assembly features, closest taxonomy annotation and relative evolutionary distance for novel genus and species.

**Supplementary Table 5:** Dereplicated MAGs with representative cluster names and their taxonomy annotations

**Supplementary Table 6:** Stats showing the diversity of GTDb taxonomy within MAGs.

**Supplementary Table 7:** Novel latin binomials for MAGs and taxa names submitted to Genbank

**Supplementary Table 8:** Comparison of genome stats between MAGs from this study and GTDb corresponding representative MAG cluster

**Supplementary Table 9:** Relative abundance, fold change and log ratio foldchange for all MAGs

**Supplementary Table 10:** Genomes depicted as early and late degraders according to the time the genomes showed a 2x fold change.

**Supplementary Table 11:** MAGs and their CAZyme profiles.

**Supplementary Table 12:** CaZymes counts for selected MAG clusters

**Supplementary Table 13:** media preparation materials, sources and quantity

**Supplementary Figure 1:** Principle Component Analysis (PCoA) showing the dynamics of the microbiome during the different time points and between the Carbohydrate treatment. PC1 and PC2 represent the percentage of variance explained by Principle Component (PC) 1 and 2.

**Supplementary Figure 2:** Changes in inverse Simpson index between time periods of the substrates.

**Supplementary figure 3: Box plots showing the dynamic shifts in read proportions for all binned MAGs after 0h, 6h, 12h and 24h fermentation in the model colon.** The box represents the interquartile range (IQR) (25^th^ and 75th percentile); the median is shown within the box. The whiskers indicate minimum and maximum Inter Quartile Range (IQR); dots represent outliers.

**Supplementary Figure 4:** Distribution of CAZy families per substrate and in all the genome

**Description of *Candidatus* Acetatifactor hominis** sp. nov.

*Candidatus* Acetatifactor hominis (ho’mi.nis. L. gen. masc. n. *hominis*, of a human being).

A bacterial species identified by metagenomic analyses. This species includes all bacteria with genomes that show ≥95% average nucleotide identity to the type genome for the species to which we have assigned the genome identifier rmaize_MAXBIN__038 and which is available via NCBI BioSample SAMN18871269. This is a new name for the alphanumeric GTDB species sp900066565. The GC content of the type genome is 47.74 % and the genome length is 3.05 Mbp.

**Description of *Candidatus* Aphodonaster** gen. nov.

*Candidatus* Aphodonaster (Aph.od.o.nas’ter. Gr. fem. n. *aphodos* dung; Gr. masc. n. *naster* an inhabitant; N.L. masc. n. *Aphodonaster* a microbe associated with faeces).

A bacterial genus identified by metagenomic analyses of human faeces. The genus includes all bacteria with genomes that show ≥60% average aminoacid identity to the genome of the type strain from the type species, *Candidatus* Aphodonaster merdae. This is a new name for the GTDB alphanumeric genus SFFH01. This genus has been assigned by GTDB-Tk v1.5.0 working on GTDB R06-RS202 reference data (Chaumeil et al., 2019; Parks et al., 2020) to the order *Christensenellales* and to the family *CAG-74*

**Description of *Candidatus* Aphodonaster intestinalis** sp. nov.

*Candidatus* Aphodonaster intestinalis (in.tes.ti.na’lis. N.L. masc. adj. *intestinalis,* pertaining to the intestines).

A bacterial species identified by metagenomic analyses. This species includes all bacteria with genomes that show ≥95% average nucleotide identity to the type genome for the species to which we have assigned the genome identifier T0_METABAT__97 and which is available via NCBI BioSample SAMN18871333. This is a new name for the alphanumeric GTDB species sp900548125. The GC content of the type genome is 55.44 % and the genome length is 2.54 Mbp.

**Description of *Candidatus* Aphodonaster merdae** sp. nov.

*Candidatus* Aphodonaster merdae (mer’dae. L. gen. fem. n. *merdae*, of faeces).

A bacterial species identified by metagenomic analyses. This species includes all bacteria with genomes that show ≥95% average nucleotide identity to the type genome for the species to which we have assigned the genome identifier pstarch_METABAT__69 and which is available via NCBI BioSample SAMN18871262. This is a new name for the alphanumeric GTDB species sp900542395. The GC content of the type genome is 59.61 % and the genome length is 2.66 Mbp.

**Description of *Candidatus* Avimicrobium caecorum** sp. nov.

*Candidatus* Avimicrobium caecorum (cae.co’rum. N. L. gen. pl. n. *caecorum*, of caeca).

A bacterial species identified by metagenomic analyses. This species includes all bacteria with genomes that show ≥95% average nucleotide identity to the type genome for the species to which we have assigned the genome identifier avicell_METABAT__34 and which is available via NCBI BioSample SAMN18871193. This is a new name for the alphanumeric GTDB species sp900547185. This genus was named by Glendinning et al. (2020). The GC content of the type genome is 56.83 % and the genome length is 2.20 Mbp.

**Description of *Candidatus* Blautia hennigii** sp. nov.

*Candidatus* Blautia hennigii (hen.ni’gi.i. N.L. gen. masc. n. *hennigii* derived from the Latinised family name for Willi Hennig, 1913-1976, the East German scientist who founded phylogenetic systematics or cladistics).

A bacterial species identified by metagenomic analyses. This species includes all bacteria with genomes that show ≥95% average nucleotide identity to the type genome for the species to which we have assigned the genome identifier hylon_METABAT__127 and which is available via NCBI BioSample SAMN18871203. This is a new name for the alphanumeric GTDB species sp900066505. GTDB has assigned this species to genus with an alphabetic suffix which cannot be incorporated into a well-formed binomial, so in naming this species, we have used the basonym for the genus. The GC content of the type genome is 43.26 % and the genome length is 2.93 Mbp.

**Description of *Candidatus* Caccadaptatus** gen. nov.

*Candidatus* Caccadaptatus (Cacc.ad.ap.ta’tus. Gr. fem. n. *kakké* dung; L. masc. part. adj. *adaptatus* adapted to; N.L. masc. n. *Caccadaptatus* a microbe associated with faeces).

A bacterial genus identified by metagenomic analyses of human faeces. The genus includes all bacteria with genomes that show ≥60% average aminoacid identity to the genome of the type strain from the type species, *Candidatus* Caccadaptatus darwinii. This is a new name for the GTDB alphanumeric genus NK3B98. This genus has been assigned by GTDB-Tk v1.5.0 working on GTDB R06-RS202 reference data (Chaumeil et al., 2019; Parks et al., 2020) to the order *Oscillospirales* and to the family *Oscillospiraceae*

**Description of *Candidatus* Caccadaptatus darwinii** sp. nov.

*Candidatus* Caccadaptatus darwinii (dar.wi’ni.i. N.L. gen. masc. n. darwinii derived from the Latinised family name for Charles Darwin, 1809-1882, the British scientist who proposed the theory of evolution by natural selection).

A bacterial species identified by metagenomic analyses. This species includes all bacteria with genomes that show ≥95% average nucleotide identity to the type genome for the species to which we have assigned the genome identifier rmaize_METABAT__56 and which is available via NCBI BioSample SAMN18871284. This is a new name for the alphanumeric GTDB species sp900545815. The GC content of the type genome is 56.11 % and the genome length is 2.31 Mbp.

**Description of *Candidatus* Chesmatocola** gen. nov.

*Candidatus* Chesmatocola (Ches.ma.to’co.la. Gr. neut. n. *chesma* dung; N.L. masc./fem. suffix *cola* an inhabitant of; N.L. fem. n. *Chesmatocola* a microbe associated with faeces).

A bacterial genus identified by metagenomic analyses of human faeces. The genus includes all bacteria with genomes that show ≥60% average aminoacid identity to the genome of the type strain from the type species, *Candidatus* Chesmatocola anthropi. This is a new name for the GTDB alphanumeric genus CAG-354. This genus has been assigned by GTDB-Tk v1.5.0 working on GTDB R06-RS202 reference data (Chaumeil et al., 2019; Parks et al., 2020) to the order *TANB77* and to the family *CAG-508*

**Description of *Candidatus* Chesmatocola anthropi** sp. nov.

*Candidatus* Chesmatocola anthropi (an.thro’pi. Gr. masc. n. *anthropos,* a human being; N.L. gen. masc. n. *anthropi,* of a human being).

A bacterial species identified by metagenomic analyses. This species includes all bacteria with genomes that show ≥95% average nucleotide identity to the type genome for the species to which we have assigned the genome identifier hylon_METABAT__79 and which is available via NCBI BioSample SAMN18871215. This is a new name for the alphanumeric GTDB species sp001915925. The GC content of the type genome is 28.31 % and the genome length is 1.38 Mbp.

**Description of *Candidatus* Cholicomonas** gen. nov.

*Candidatus* Cholicomonas (Cho.li.co.mo’nas. Gr. fem. n. *cholix, cholikos* guts; L. fem. n. *monas* a monad; N.L. fem. n. *Cholicomonas* a microbe associated with the intestines).

A bacterial genus identified by metagenomic analyses of human faeces. The genus includes all bacteria with genomes that show ≥60% average aminoacid identity to the genome of the type strain from the type species, *Candidatus* Cholicomonas copri. This is a new name for the GTDB alphanumeric genus CAG-628. This genus has been assigned by GTDB-Tk v1.5.0 working on GTDB R06-RS202 reference data (Chaumeil et al., 2019; Parks et al., 2020) to the order *RF39* and to the family *UBA660*

**Description of *Candidatus* Cholicomonas copri** sp. nov.

*Candidatus* Cholicomonas copri (cop’ri. Gr. masc. n. *kópros*, faeces; N.L. gen. n. copri; of faeces).

A bacterial species identified by metagenomic analyses. This species includes all bacteria with genomes that show ≥95% average nucleotide identity to the type genome for the species to which we have assigned the genome identifier T0_METABAT__30 and which is available via NCBI BioSample SAMN18871328. This is a new name for the alphanumeric GTDB species sp000438415. The GC content of the type genome is 27.35 % and the genome length is 0.62 Mbp.

**Description of *Candidatus* Choliconaster** gen. nov.

*Candidatus* Choliconaster (Cho.li.co.nas’ter. Gr. fem. n. *cholix, cholikos*guts; Gr. masc. n. *naster* an inhabitant; N.L. masc. n. *Choliconaster* a microbe associated with the intestines).

A bacterial genus identified by metagenomic analyses of human faeces. The genus includes all bacteria with genomes that show ≥60% average aminoacid identity to the genome of the type strain from the type species, *Candidatus* Choliconaster caccae. This is a new name for the GTDB alphanumeric genus ER4. This genus has been assigned by GTDB-Tk v1.5.0 working on GTDB R06-RS202 reference data (Chaumeil et al., 2019; Parks et al., 2020) to the order *Oscillospirales* and to the family *Oscillospiraceae*

**Description of *Candidatus* Choliconaster caccae** sp. nov.

*Candidatus* Choliconaster caccae (cac’cae. Gr. fem. n. *kakkê*, faeces; N.L. gen. n. *caccae*, of faeces).

A bacterial species identified by metagenomic analyses. This species includes all bacteria with genomes that show ≥95% average nucleotide identity to the type genome for the species to which we have assigned the genome identifier T0_MAXBIN__028 and which is available via NCBI BioSample SAMN18871294. This is a new name for the alphanumeric GTDB species sp000765235. The GC content of the type genome is 57.68 % and the genome length is 2.86 Mbp.

**Description of *Candidatus* Choliconaster merdae** sp. nov.

*Candidatus* Choliconaster merdae (mer’dae. L. gen. fem. n. *merdae*, of faeces).

A bacterial species identified by metagenomic analyses. This species includes all bacteria with genomes that show ≥95% average nucleotide identity to the type genome for the species to which we have assigned the genome identifier T0_METABAT__240 and which is available via NCBI BioSample SAMN18871322. This is a new name for the alphanumeric GTDB species sp900317525. The GC content of the type genome is 60.79 % and the genome length is 1.92 Mbp.

**Description of *Candidatus* Clostridium faecihominis** sp. nov.

*Candidatus* Clostridium faecihominis (fae.ci.ho’mi.nis. L. fem. n. *faex, faecis* faeces; L. gen. masc. n. *hominis*, of a human being; N.L. gen. n. *faecihominis*, of human faeces).

A bacterial species identified by metagenomic analyses. This species includes all bacteria with genomes that show ≥95% average nucleotide identity to the type genome for the species to which we have assigned the genome identifier T0_METABAT__195 and which is available via NCBI BioSample SAMN18871315. This is a new name for the alphanumeric GTDB species sp003024715. GTDB has assigned this species to genus with an alphabetic suffix which cannot be incorporated into a well-formed binomial, so in naming this species, we have used the basonym for the genus. The GC content of the type genome is 49.01 % and the genome length is 2.60 Mbp.

**Description of *Candidatus* Colibacterium** gen. nov.

*Candidatus* Colibacterium (Co.li.bac.te’ri.um. L. neut. n. *colon* large intestine; N.L. neut. n. *bacterium* a bacterium; N.L. neut. n. *Colibacterium* a microbe associated with the intestines).

A bacterial genus identified by metagenomic analyses of human faeces. The genus includes all bacteria with genomes that show ≥60% average aminoacid identity to the genome of the type strain from the type species, *Candidatus* Colibacterium hominis. This is a new name for the GTDB alphanumeric genus SFEL01. This genus has been assigned by GTDB-Tk v1.5.0 working on GTDB R06-RS202 reference data (Chaumeil et al., 2019; Parks et al., 2020) to the order *Christensenellales* and to the family *CAG-138*

**Description of *Candidatus* Colibacterium hominis** sp. nov.

*Candidatus* Colibacterium hominis (ho’mi.nis. L. gen. masc. n. *hominis*, of a human being).

A bacterial species identified by metagenomic analyses. This species includes all bacteria with genomes that show ≥95% average nucleotide identity to the type genome for the species to which we have assigned the genome identifier T0_MAXBIN__134 and which is available via NCBI BioSample SAMN18871298. This is a new name for the alphanumeric GTDB species sp004557245. The GC content of the type genome is 54.28 % and the genome length is 1.56 Mbp.

**Description of *Candidatus* Colihabitans** gen. nov.

*Candidatus* Colihabitans (Co.li.ha’bi.tans. L. neut. n. *colon* large intestine; L. masc./fem. adj. part. *habitans* an inhabitant; N.L. fem. n. *Colihabitans* a microbe associated with the intestines).

A bacterial genus identified by metagenomic analyses of human faeces. The genus includes all bacteria with genomes that show ≥60% average aminoacid identity to the genome of the type strain from the type species, *Candidatus* Colihabitans norwichensis. This is a new name for the GTDB alphanumeric genus CAG-170. This genus has been assigned by GTDB-Tk v1.5.0 working on GTDB R06-RS202 reference data (Chaumeil et al., 2019; Parks et al., 2020) to the order *Oscillospirales* and to the family *Oscillospiraceae*

**Description of *Candidatus* Colihabitans hominis** sp. nov.

*Candidatus* Colihabitans hominis (ho’mi.nis. L. gen. masc. n. *hominis*, of a human being).

A bacterial species identified by metagenomic analyses. This species includes all bacteria with genomes that show ≥95% average nucleotide identity to the type genome for the species to which we have assigned the genome identifier T0_METABAT__180 and which is available via NCBI BioSample SAMN18871312. This is a new name for the alphanumeric GTDB species sp900549635. The GC content of the type genome is 56.47 % and the genome length is 2.50 Mbp.

**Description of *Candidatus* Colihabitans norwichensis** sp. nov.

*Candidatus* Colihabitans norwichensis (nor.wich.en’sis. N.L. fem. adj. *norwichensis* pertaining to English city of Norwich).

A bacterial species identified by metagenomic analyses. This species includes all bacteria with genomes that show ≥95% average nucleotide identity to the type genome for the species to which we have assigned the genome identifier hylon_METABAT__172 and which is available via NCBI BioSample SAMN18871205. This is a new name for the alphanumeric GTDB species sp000432135. The GC content of the type genome is 57.56 % and the genome length is 3.16 Mbp.

**Description of *Candidatus* Dysosmobacter stercoris** sp. nov.

*Candidatus* Dysosmobacter stercoris (ster’co.ris. L. gen. neut. n. *stercoris*, of faeces).

A bacterial species identified by metagenomic analyses. This species includes all bacteria with genomes that show ≥95% average nucleotide identity to the type genome for the species to which we have assigned the genome identifier hylon_METABAT__95 and which is available via NCBI BioSample SAMN18871217. This is a new name for the alphanumeric GTDB species sp900542115. The GC content of the type genome is 58.43 % and the genome length is 1.44 Mbp.

**Description of *Candidatus* Eisenbergiella faecalis** sp. nov.

*Candidatus* Eisenbergiella faecalis (fae.ca’lis. N.L. fem. adj. *faecalis*, of faeces).

A bacterial species identified by metagenomic analyses. This species includes all bacteria with genomes that show ≥95% average nucleotide identity to the type genome for the species to which we have assigned the genome identifier pstarch_METABAT__22 and which is available via NCBI BioSample SAMN18871259. This is a new name for the alphanumeric GTDB species sp900066775. The GC content of the type genome is 48.80 % and the genome length is 2.82 Mbp.

**Description of *Candidatus* Enteromorpha** gen. nov.

*Candidatus* Enteromorpha (En.te.ro.mor’pha. Gr. neut. n. *enteron* the gut; Gr. fem. n. *morphe* a form, shape; N.L. fem. n. *Enteromorpha* a microbe associated with the intestines).

A bacterial genus identified by metagenomic analyses of human faeces. The genus includes all bacteria with genomes that show ≥60% average aminoacid identity to the genome of the type strain from the type species, *Candidatus* Enteromorpha quadrami. This is a new name for the GTDB alphanumeric genus CAG-110. This genus has been assigned by GTDB-Tk v1.5.0 working on GTDB R06-RS202 reference data (Chaumeil et al., 2019; Parks et al., 2020) to the order *Oscillospirales* and to the family *Oscillospiraceae*

**Description of *Candidatus* Enteromorpha barnesiae** sp. nov.

*Candidatus* Enteromorpha barnesiae (bar.ne’si.ae. N.L. gen. fem. n. *barnesiae*, of Barnes, named after Ella M. Barnes, a British microbiologist).

A bacterial species identified by metagenomic analyses. This species includes all bacteria with genomes that show ≥95% average nucleotide identity to the type genome for the species to which we have assigned the genome identifier avicell_METABAT__70, hylon_METABAT__9, inulin_METABAT__82 and rmaize_METABAT__177 and which is available via NCBI BioSample SAMN18871197. This is a new name for the alphanumeric GTDB species sp003525905. The GC content of the type genome is 61.70 %, 61.32 %, 61.45 % and 62.01 % and the genome length is 1.70 Mbp, 2.26 Mbp, 2.12 Mbp and 1.81 Mbp.

**Description of *Candidatus* Enteromorpha quadrami** sp. nov.

*Candidatus* Enteromorpha quadrami (quad.ra’mi. N.L. gen. n. *quadrami* of the Quadram Institute).

A bacterial species identified by metagenomic analyses. This species includes all bacteria with genomes that show ≥95% average nucleotide identity to the type genome for the species to which we have assigned the genome identifiers avicel_MAXBIN__045, inulin_METABAT__94 and pstarch_METABAT__151 and which is available via NCBI BioSample SAMN18871185. This is a new name for the alphanumeric GTDB species sp000434635. The GC content of the type genome is 57.48 %, 57.06 % and 57.30 5 and the genome length are 2.09 Mbp, 2.31 Mbp and 2.27 Mbp.

**Description of *Candidatus* Enteronaster** gen. nov.

*Candidatus* Enteronaster (En.ter.o.nas’ter. Gr. neut. n. *enteron* the gut; Gr. masc. n. *naster* an inhabitant; N.L. masc. n. *Enteronaster* a microbe associated with the intestines).

A bacterial genus identified by metagenomic analyses of human faeces. The genus includes all bacteria with genomes that show ≥60% average aminoacid identity to the genome of the type strain from the type species, *Candidatus* Enteronaster faecalis. This is a new name for the GTDB alphanumeric genus CAG-103. This genus has been assigned by GTDB-Tk v1.5.0 working on GTDB R06-RS202 reference data (Chaumeil et al., 2019; Parks et al., 2020) to the order *Oscillospirales* and to the family *Oscillospiraceae*

**Description of *Candidatus* Enteronaster faecalis** sp. nov.

*Candidatus* Enteronaster faecalis (fae.ca’lis. N.L. masc. adj. *faecalis*, of faeces).

A bacterial species identified by metagenomic analyses. This species includes all bacteria with genomes that show ≥95% average nucleotide identity to the type genome for the species to which we have assigned the genome identifier inulin_METABAT__98 and which is available via NCBI BioSample SAMN18871242. This is a new name for the alphanumeric GTDB species sp000432375. The GC content of the type genome is 61.98 % and the genome length is 1.97 Mbp.

**Description of *Candidatus* Enteroplasma** gen. nov.

*Candidatus* Enteroplasma (En.te.ro.plas’ma. Gr. neut. n. *enteron* the gut; L. neut. n. *plasma* a form; N.L. neut. n. *Enteroplasma* a microbe associated with the intestines).

A bacterial genus identified by metagenomic analyses of human faeces. The genus includes all bacteria with genomes that show ≥60% average aminoacid identity to the genome of the type strain from the type species, *Candidatus* Enteroplasma stercoris. This is a new name for the GTDB alphanumeric genus CAG-115. This genus has been assigned by GTDB-Tk v1.5.0 working on GTDB R06-RS202 reference data (Chaumeil et al., 2019; Parks et al., 2020) to the order *Oscillospirales* and to the family *Ruminococcaceae*

**Description of *Candidatus* Enteroplasma stercoris** sp. nov.

*Candidatus* Enteroplasma stercoris (ster’co.ris. L. gen. neut. n. *stercoris*, of faeces).

A bacterial species identified by metagenomic analyses. This species includes all bacteria with genomes that show ≥95% average nucleotide identity to the type genome for the species to which we have assigned the genome identifier inulin_MAXBIN__035 and which is available via NCBI BioSample SAMN18871220. This is a new name for the alphanumeric GTDB species sp003531585. The GC content of the type genome is 52.91 % and the genome length is 2.79 Mbp.

**Description of *Candidatus* Enterovivens** gen. nov.

*Candidatus* Enterovivens (En.te.ro.vi’vens. Gr. neut. n. *enteron* the gut; N.L. masc./fem. adj. part. *vivens* living; N.L. fem. n. *Enterovivens* a microbe living in the intestines).

A bacterial genus identified by metagenomic analyses of human faeces. The genus includes all bacteria with genomes that show ≥60% average aminoacid identity to the genome of the type strain from the type species, *Candidatus* Enterovivens caccae. This is a new name for the GTDB alphanumeric genus CAG-127. This genus has been assigned by GTDB-Tk v1.5.0 working on GTDB R06-RS202 reference data (Chaumeil et al., 2019; Parks et al., 2020) to the order *Lachnospirales* and to the family *Lachnospiraceae*

**Description of *Candidatus* Enterovivens caccae** sp. nov.

*Candidatus* Enterovivens caccae (cac’cae. Gr. fem. n. *kakkê*, faeces; N.L. gen. n. *caccae*, of faeces).

A bacterial species identified by metagenomic analyses. This species includes all bacteria with genomes that show ≥95% average nucleotide identity to the type genome for the species to which we have assigned the genome identifier T0_METABAT__183 and which is available via NCBI BioSample SAMN18871313. This is a new name for the alphanumeric GTDB species sp900319515. The GC content of the type genome is 44.48 % and the genome length is 2.61 Mbp.

**Description of *Candidatus* Eubacterium caccanthorpi** sp. nov.

*Candidatus* Eubacterium caccanthorpi (cacc.an.thro’pi. Gr. fem. n. *kakkê*, faeces; Gr. masc. n. *anthropos*, a human being; N.L. gen. masc. n. *caccanthropi*, of human faeces).

A bacterial species identified by metagenomic analyses. This species includes all bacteria with genomes that show ≥95% average nucleotide identity to the type genome for the species to which we have assigned the genome identifier T0_MAXBIN__022 and which is available via NCBI BioSample SAMN18871293. This is a new name for the alphanumeric GTDB species sp000434995. GTDB has assigned this species to genus with an alphabetic suffix which cannot be incorporated into a well-formed binomial, so in naming this species, we have used the basonym for the genus. The GC content of the type genome is 36.52 % and the genome length is 1.94 Mbp.

**Description of *Candidatus* Eubacterium colihabitans** sp. nov.

*Candidatus* Eubacterium colihabitans (co.li.ha’bi.tans. L. neut. n. *colum*, colon; L. pres. part. *habitans*, inhabiting; N.L. part. adj. *colihabitans*, inhabiting the colon).

A bacterial species identified by metagenomic analyses. This species includes all bacteria with genomes that show ≥95% average nucleotide identity to the type genome for the species to which we have assigned the genome identifier T0_METABAT__220 and which is available via NCBI BioSample SAMN18871319. This is a new name for the alphanumeric GTDB species sp003491505. GTDB has assigned this species to genus with an alphabetic suffix which cannot be incorporated into a well-formed binomial, so in naming this species, we have used the basonym for the genus. The GC content of the type genome is 41.07 % and the genome length is 2.49 Mbp.

**Description of *Candidatus* Gallacutalibacter hominis** sp. nov.

*Candidatus* Gallacutalibacter hominis (ho’mi.nis. L. gen. masc. n. *hominis*, of a human being).

A bacterial species identified by metagenomic analyses. This species includes all bacteria with genomes that show ≥95% average nucleotide identity to the type genome for the species to which we have assigned the genome identifiers avicel METABAT__20 and inulin_METABAT__180 and which is available via NCBI BioSample SAMN18871192. This is a new name for the alphanumeric GTDB species sp003477405. This genus was named by Gilroy et al. (2021). The GC content of the type genomes is 56.15 % and 56. 25 % and the genome lengths are 2.33 Mbp and 1.92 Mbp.

**Description of *Candidatus* Gemmiger merdicola** sp. nov.

*Candidatus* Gemmiger merdicola (mer.di’co.la. L. gen. fem. n. *merda*, faeces; L. masc./fem. suff. *-cola*, inhabitant of; N.L. fem. n. *merdicola* inhabitant of faeces).

A bacterial species identified by metagenomic analyses. This species includes all bacteria with genomes that show ≥95% average nucleotide identity to the type genome for the species to which we have assigned the genome identifier inulin_METABAT__93 and which is available via NCBI BioSample SAMN18871240. This is a new name for the alphanumeric GTDB species sp900539695. The GC content of the type genome is 58.43 % and the genome length is 2.39 Mbp.

**Description of *Candidatus* Holdemanella enterica** sp. nov.

*Candidatus* Holdemanella enterica (en.ter’i.ca. Gr. neut. n. *enteron*, gut; L. fem. adj. suff. *-ica*, pertaining to; N.L. fem. adj.*enterica*, pertaining to the gut).

A bacterial species identified by metagenomic analyses. This species includes all bacteria with genomes that show ≥95% average nucleotide identity to the type genome for the species to which we have assigned the genome identifier T0_METABAT__130 and which is available via NCBI BioSample SAMN18871306. This is a new name for the alphanumeric GTDB species sp002299315. The GC content of the type genome is 34.07 % and the genome length is 2.18 Mbp.

**Description of *Candidatus* Huxleyella** gen. nov.

*Candidatus* Huxleyella (Hux.ley.el’la. L. fem. dim. suff. *-ella* diminutive ending; N.L. fem. n. *Huxleyella* named in honour of the British scientist Thomas Henry Huxley (1825-1895), known for his advocacy of Charles Darwin’s theory of evolution).

A bacterial genus identified by metagenomic analyses of human faeces. The genus includes all bacteria with genomes that show ≥60% average aminoacid identity to the genome of the type strain from the type species, *Candidatus* Huxleyella fimi. This is a new name for the GTDB alphanumeric genus UMGS1071. This genus has been assigned by GTDB-Tk v1.5.0 working on GTDB R06-RS202 reference data (Chaumeil et al., 2019; Parks et al., 2020) to the order *Oscillospirales* and to the family *Acutalibacteraceae*

**Description of *Candidatus* Huxleyella fimi** sp. nov.

*Candidatus* Huxleyella fimi (fi’mi. L. neut. gen. n. *fimi*, of faeces).

A bacterial species identified by metagenomic analyses. This species includes all bacteria with genomes that show ≥95% average nucleotide identity to the type genome for the species to which we have assigned the genome identifier T0_METABAT__173 and which is available via NCBI BioSample SAMN18871311. This is a new name for the alphanumeric GTDB species sp900542375. The GC content of the type genome is 38.84 % and the genome length is 1.60 Mbp.

**Description of *Candidatus* Minthomorpha** gen. nov.

*Candidatus* Minthomorpha (Min.tho.mor’pha. Gr. masc. n. *minthos* dung; Gr. fem. n. *morphe* a form, shape; N.L. fem. n. *Minthomorpha* a microbe associated with faeces).

A bacterial genus identified by metagenomic analyses of human faeces. The genus includes all bacteria with genomes that show ≥60% average aminoacid identity to the genome of the type strain from the type species, *Candidatus* Minthomorpha faecalis. This is a new name for the GTDB alphanumeric genus CAG-81. This genus has been assigned by GTDB-Tk v1.5.0 working on GTDB R06-RS202 reference data (Chaumeil et al., 2019; Parks et al., 2020) to the order *Lachnospirales* and to the family *Lachnospiraceae*

**Description of *Candidatus* Minthomorpha faecalis** sp. nov.

*Candidatus* Minthomorpha faecalis (fae.ca’lis. N.L. fem. adj. *faecalis*, of faeces).

A bacterial species identified by metagenomic analyses. This species includes all bacteria with genomes that show ≥95% average nucleotide identity to the type genome for the species to which we have assigned the genome identifier rmaize_METABAT__174 and which is available via NCBI BioSample SAMN18871281. This is a new name for the alphanumeric GTDB species sp900066535. The GC content of the type genome is 49.05 % and the genome length is 2.98 Mbp.

**Description of *Candidatus* Minthonaster** gen. nov.

*Candidatus* Minthonaster (Min.tho.nas’ter. Gr. masc. n. *minthos* dung; Gr. masc. n. *naster* an inhabitant; N.L. masc. n. *Minthonaster* a microbe associated with faeces).

A bacterial genus identified by metagenomic analyses of human faeces. The genus includes all bacteria with genomes that show ≥60% average aminoacid identity to the genome of the type strain from the type species, *Candidatus* Minthonaster faecium. This is a new name for the GTDB alphanumeric genus CAG-83. This genus has been assigned by GTDB-Tk v1.5.0 working on GTDB R06-RS202 reference data (Chaumeil et al., 2019; Parks et al., 2020) to the order *Oscillospirales* and to the family *Oscillospiraceae*

**Description of *Candidatus* Minthonaster anthropi** sp. nov.

*Candidatus* Minthonaster anthropi (an.thro’pi. Gr. masc. n. *anthropos,* a human being; N.L. gen. masc. n. *anthropi*, of a human being).

A bacterial species identified by metagenomic analyses. This species includes all bacteria with genomes that show ≥95% average nucleotide identity to the type genome for the species to which we have assigned the genome identifier T0_METABAT__167 and which is available via NCBI BioSample SAMN18871310. This is a new name for the alphanumeric GTDB species sp900552475. The GC content of the type genome is 61.38 % and the genome length is 2.18 Mbp.

**Description of *Candidatus* Minthonaster faecium** sp. nov.

*Candidatus* Minthonaster faecium (fae’ci.um. L. fem. gen. pl. n. *faecium,* of faeces).

A bacterial species identified by metagenomic analyses. This species includes all bacteria with genomes that show ≥95% average nucleotide identity to the type genome for the species to which we have assigned the genome identifier hylon_METABAT__44 and which is available via NCBI BioSample SAMN18871213. This is a new name for the alphanumeric GTDB species sp003539495. The GC content of the type genome is 57.03 % and the genome length is 2.06 Mbp.

**Description of *Candidatus* Minthonaster hominis** sp. nov.

*Candidatus* Minthonaster hominis (ho’mi.nis. L. gen. masc. n. *hominis*, of a human being).

A bacterial species identified by metagenomic analyses. This species includes all bacteria with genomes that show ≥95% average nucleotide identity to the type genome for the species to which we have assigned the genome identifier inulin_METABAT__175 and which is available via NCBI BioSample SAMN18871228. This is a new name for the alphanumeric GTDB species sp900545585. The GC content of the type genome is 60.55 % and the genome length is 2.24 Mbp.

**Description of *Candidatus* Minthonaster merdae** sp. nov.

*Candidatus* Minthonaster merdae (mer’dae. L. gen. fem. n. *merdae*, of faeces).

A bacterial species identified by metagenomic analyses. This species includes all bacteria with genomes that show ≥95% average nucleotide identity to the type genome for the species to which we have assigned the genome identifier T0_METABAT__66 and which is available via NCBI BioSample SAMN18871332. This is a new name for the alphanumeric GTDB species sp000431575. The GC content of the type genome is 59.89 % and the genome length is 2.00 Mbp.

**Description of *Candidatus* Minthoplasma** gen. nov.

*Candidatus* Minthoplasma (Min.tho.plas’ma. Gr. masc. n. *minthos* dung; L. neut. n. *plasma* a form; *Minthoplasma* a microbe associated with faeces).

A bacterial genus identified by metagenomic analyses of human faeces. The genus includes all bacteria with genomes that show ≥60% average aminoacid identity to the genome of the type strain from the type species, *Candidatus* Minthoplasma entericum. This is a new name for the GTDB alphanumeric genus GCA-900066135. This genus has been assigned by GTDB-Tk v1.5.0 working on GTDB R06-RS202 reference data (Chaumeil et al., 2019; Parks et al., 2020) to the order *Lachnospirales* and to the family *Lachnospiraceae*

**Description of *Candidatus* Minthoplasma copri** sp. nov.

*Candidatus* Minthoplasma copri (cop’ri. Gr. masc. n. kópros, faeces; N.L. gen. n. copri; of faeces).

A bacterial species identified by metagenomic analyses. This species includes all bacteria with genomes that show ≥95% average nucleotide identity to the type genome for the species to which we have assigned the genome identifier T0_METABAT__206 and which is available via NCBI BioSample SAMN18871318. This is a new name for the alphanumeric GTDB species sp900543575. The GC content of the type genome is 49.81 % and the genome length is 3.26 Mbp.

**Description of *Candidatus* Minthoplasma enterica** sp. nov.

*Candidatus* Minthoplasma entericum (en.te’ri.cum. Gr. neut. n. *enteron*, gut; L. neut. adj. suff. *-icum*, pertaining to; N.L. neut. adj.*entericum*, pertaining to the gut).

A bacterial species identified by metagenomic analyses. This species includes all bacteria with genomes that show ≥95% average nucleotide identity to the type genome for the species to which we have assigned the genome identifier T0_METABAT__122 and which is available via NCBI BioSample SAMN18871303. This is a new name for the alphanumeric GTDB species sp900066135. The GC content of the type genome is 47.02 % and the genome length is 1.90 Mbp.

**Description of *Candidatus* Minthovivens** gen. nov.

*Candidatus* Minthovivens (Min.tho.viv’ens. Gr. masc. n. *minthos* dung; N.L. masc./fem. part. adj. *vivens* living; N.L. fem. n. *Minthovivens* a microbe living in faeces).

A bacterial genus identified by metagenomic analyses of human faeces. The genus includes all bacteria with genomes that show ≥60% average aminoacid identity to the genome of the type strain from the type species, *Candidatus* Minthovivens enterohominis. This is a new name for the GTDB alphanumeric genus KLE1615. This genus has been assigned by GTDB-Tk v1.5.0 working on GTDB R06-RS202 reference data (Chaumeil et al., 2019; Parks et al., 2020) to the order *Lachnospirales* and to the family *Lachnospiraceae*

**Description of *Candidatus* Minthovivens enterohominis** sp. nov.

*Candidatus* Minthovivens enterohominis (en.te.ro.ho’mi.nis. Gr. neut. n. *enteron*, gut; L. gen. masc. n. *hominis,* of a human being; N.L. gen. masc. n. *enterohominis*, of the human gut).

A bacterial species identified by metagenomic analyses. This species includes all bacteria with genomes that show ≥95% average nucleotide identity to the type genome for the species to which we have assigned the genome identifier inulin_METABAT__130 and which is available via NCBI BioSample SAMN18871226. This is a new name for the alphanumeric GTDB species sp900066985. The GC content of the type genome is 40.97 % and the genome length is 3.77 Mbp.

**Description of *Candidatus* Negativibacillus quadrami** sp. nov.

*Candidatus* Negativibacillus quadrami (quad.ra’mi. N.L. gen. n. *quadrami* of the Quadram Institute).

A bacterial species identified by metagenomic analyses. This species includes all bacteria with genomes that show ≥95% average nucleotide identity to the type genome for the species to which we have assigned the genome identifier T0_METABAT__114 and which is available via NCBI BioSample SAMN18871301. This is a new name for the alphanumeric GTDB species sp000435195. The GC content of the type genome is 51.95 % and the genome length is 2.20 Mbp.

**Description of *Candidatus* Neoacutalibacter** gen. nov.

*Candidatus* Neoacutalibacter (Ne.o.a.cu.ta.li.ibac’ter. Gr. masc. adj. *neos* new; N.L. masc. n. *Acutalibacter* an existing genus name; N.L. masc. n. *Neoacutalibacter* a bacterial genus related to but distinct from the existing named genus).

A bacterial genus identified by metagenomic analyses of human faeces. The genus includes all bacteria with genomes that show ≥60% average aminoacid identity to the genome of the type strain from the type species, *Candidatus* Neoacutalibacter hominis. This is a new name for the GTDB alphanumeric genus CAG-177. This genus has been assigned by GTDB-Tk v1.5.0 working on GTDB R06-RS202 reference data (Chaumeil et al., 2019; Parks et al., 2020) to the order *Oscillospirales* and to the family *Acutalibacteraceae*

**Description of *Candidatus* Neoacutalibacter hominis** sp. nov.

*Candidatus* Neoacutalibacter hominis (ho’mi.nis. L. gen. masc. n. *hominis*, of a human being).

A bacterial species identified by metagenomic analyses. This species includes all bacteria with genomes that show ≥95% average nucleotide identity to the type genome for the species to which we have assigned the genome identifier inulin_MAXBIN__022 and which is available via NCBI BioSample SAMN18871219. This is a new name for the alphanumeric GTDB species sp003514385. The GC content of the type genome is 51.47 % and the genome length is 2.22 Mbp.

**Description of *Candidatus* Neoanaerovorax** gen. nov.

*Candidatus* Neoanaerovorax (Ne.o.an.ae.ro.vo’rax. Gr. masc. adj. *neos* new; N.L. masc. n. *Anaerovorax* an existing genus name; N.L. masc. n. *Neoanaerovorax* a bacterial genus related to but distinct from the existing named genus).

A bacterial genus identified by metagenomic analyses of human faeces. The genus includes all bacteria with genomes that show ≥60% average aminoacid identity to the genome of the type strain from the type species, *Candidatus* Neoanaetovorax merdae. This is a new name for the GTDB alphanumeric genus CAG-238. This genus has been assigned by GTDB-Tk v1.5.0 working on GTDB R06-RS202 reference data (Chaumeil et al., 2019; Parks et al., 2020) to the order *Peptostreptococcales* and to the family *Anaerovoracaceae*

**Description of *Candidatus* Neoanaerovorax merdae** sp. nov.

*Candidatus* Neoanaerovorax merdae (mer’dae. L. gen. fem. n. *merdae*, of faeces).

A bacterial species identified by metagenomic analyses. This species includes all bacteria with genomes that show ≥95% average nucleotide identity to the type genome for the species to which we have assigned the genome identifiers rmaize_METABAT__46_sub and T0_METABAT__161 and which is available via NCBI BioSample SAMN18871283. This is a new name for the alphanumeric GTDB species sp900542245. The GC content of the type genome are 52.09 % and 51.52 % and the genome lengths are 1.57 Mbp and 2.01 Mbp.

**Description of *Candidatus* Neoeggerthella** gen. nov.

*Candidatus* Neoeggerthella (Ne.o.eg.ger.thel’la. Gr. masc. adj. *neos* new; N.L. fem. n. *Eggerthella* an existing genus name; N.L. fem. n. *Neoeggerthella* a bacterial genus related to but distinct from the existing named genus).

A bacterial genus identified by metagenomic analyses of human faeces. The genus includes all bacteria with genomes that show ≥60% average aminoacid identity to the genome of the type strain from the type species, *Candidatus* Neoeggerthella hominis. This is a new name for the GTDB alphanumeric genus CAG-1427. This genus has been assigned by GTDB-Tk v1.5.0 working on GTDB R06-RS202 reference data (Chaumeil et al., 2019; Parks et al., 2020) to the order *Coriobacteriales* and to the family *Eggerthellaceae*

**Description of *Candidatus* Neoeggerthella hominis** sp. nov.

*Candidatus* Neoeggerthella hominis (ho’mi.nis. L. gen. masc. n. *hominis*, of a human being).

A bacterial species identified by metagenomic analyses. This species includes all bacteria with genomes that show ≥95% average nucleotide identity to the type genome for the species to which we have assigned the genome identifier nmaize_METABAT__52 and which is available via NCBI BioSample SAMN18871250. This is a new name for the alphanumeric GTDB species sp900554685. The GC content of the type genome is 45.89 % and the genome length is 1.92 Mbp.

**Description of *Candidatus* Pararuminococcus** gen. nov.

*Candidatus* Pararuminococcus (Pa.ra.ru.mi.no.coc’cus. Gr. pref. *para-*beside; N.L. masc. n. *Ruminococcus* an existing genus name; N.L. masc. n. *Pararuminococcus* a bacterial genus related to but distinct from the existing named genus).

A bacterial genus identified by metagenomic analyses of human faeces. The genus includes all bacteria with genomes that show ≥60% average aminoacid identity to the genome of the type strain from the type species, *Candidatus* Parauminococcus sangeri. This is a new name for the GTDB alphanumeric genus UBA1417. This genus has been assigned by GTDB-Tk v1.5.0 working on GTDB R06-RS202 reference data (Chaumeil et al., 2019; Parks et al., 2020) to the order *Oscillospirales* and to the family *Acutalibacteraceae*

**Description of *Candidatus* Parasutterella caccanthropi** sp. nov.

*Candidatus* Parasutterella caccanthropi (cacc.an.thro’pi. Gr. fem. n. *kakkê*, faeces; Gr. masc. n. *anthropos*, a human being; N.L. gen. masc. n. *caccanthropi*, of human faeces).

A bacterial species identified by metagenomic analyses. This species includes all bacteria with genomes that show ≥95% average nucleotide identity to the type genome for the species to which we have assigned the genome identifier pstarch_METABAT__57 and which is available via NCBI BioSample SAMN18871261. This is a new name for the alphanumeric GTDB species sp000980495. The GC content of the type genome is 49.32 % and the genome length is 2.19 Mbp.

**Description of *Candidatus* Parauminococcus sangeri** sp. nov.

*Candidatus* Parauminococcus sangeri (san’ge.ri. N.L. masc. n. *sangeri* derived from the Latinised family name for Frederick Sanger, 1918-2013, the British scientist; awarded the 1958 Nobel Prize in Chemistry for his work on the structure of protein and the 1980 Nobel Prize in Chemistry for inventing dideoxy sequencing).

A bacterial species identified by metagenomic analyses. This species includes all bacteria with genomes that show ≥95% average nucleotide identity to the type genome for the species to which we have assigned the genome identifier T0_METABAT__250 and which is available via NCBI BioSample SAMN18871325. This is a new name for the alphanumeric GTDB species sp003531055. The GC content of the type genome is 53.40 % and the genome length is 2.30 Mbp.

**Description of *Candidatus* Pearsonella** gen. nov.

*Candidatus* Pearsonella (Pear.son.el’la. L. fem. dim. suff. *-ella* diminutive ending; N.L. fem. n. *Pearsonella* named in honour of the British scientist Bruce Pearson, known for his contributions to the study of *Campylobacter*).

A bacterial genus identified by metagenomic analyses of human faeces. The genus includes all bacteria with genomes that show ≥60% average aminoacid identity to the genome of the type strain from the type species, *Candidatus* Pearsonella faecalis. This is a new name for the GTDB alphanumeric genus UBA1822. This genus has been assigned by GTDB-Tk v1.5.0 working on GTDB R06-RS202 reference data (Chaumeil et al., 2019; Parks et al., 2020) to the order *Veillonellales* and to the family *Dialisteraceae*

**Description of *Candidatus* Pearsonella faecalis** sp. nov.

*Candidatus* Pearsonella faecalis (fae.ca’lis. N.L. fem. adj. *faecalis*, of faeces).

A bacterial species identified by metagenomic analyses. This species includes all bacteria with genomes that show ≥95% average nucleotide identity to the type genome for the species to which we have assigned the genome identifier hylon_MAXBIN__006 and which is available via NCBI BioSample SAMN18871199. This is a new name for the alphanumeric GTDB species sp002314995. The GC content of the type genome is 56.55 % and the genome length is 1.81 Mbp.

**Description of *Candidatus* Physcomorpha** gen. nov.

*Candidatus* Physcomorpha (Phys.co.mor’pha. Gr. fem. n. *physke* large intestine; Gr. fem. n. *morphe* a form, shape; N.L. fem. n. *Physcomorpha* a microbe associated with the large intestine).

A bacterial genus identified by metagenomic analyses of human faeces. The genus includes all bacteria with genomes that show ≥60% average aminoacid identity to the genome of the type strain from the type species, *Candidatus* Physcomorpha faecium. This is a new name for the GTDB alphanumeric genus UBA11524. This genus has been assigned by GTDB-Tk v1.5.0 working on GTDB R06-RS202 reference data (Chaumeil et al., 2019; Parks et al., 2020) to the order *Christensenellales* and to the family *CAG-74*

**Description of *Candidatus* Physcomorpha faecium** sp. nov.

*Candidatus* Physcomorpha faecium (fae’ci.um. L. fem. gen. pl. n. *faecium*, of faeces).

A bacterial species identified by metagenomic analyses. This species includes all bacteria with genomes that show ≥95% average nucleotide identity to the type genome for the species to which we have assigned the genome identifier rmaize_MAXBIN__031 and which is available via NCBI BioSample SAMN18871268. This is a new name for the alphanumeric GTDB species sp000437595. The GC content of the type genome is 57.78 % and the genome length is 3.22 Mbp.

**Description of *Candidatus* Physconaster** gen. nov.

*Candidatus* Physconaster (Phys.co.nas’ter. Gr. fem. n. *physke* large intestine; Gr. masc. n. *naster* an inhabitant N.L. masc. n. *Physconaster* a microbe inhabiting the large intestine).

A bacterial genus identified by metagenomic analyses of human faeces. The genus includes all bacteria with genomes that show ≥60% average aminoacid identity to the genome of the type strain from the type species, *Candidatus* Physconaster merdicola. This is a new name for the GTDB alphanumeric genus UBA11774. This genus has been assigned by GTDB-Tk v1.5.0 working on GTDB R06-RS202 reference data (Chaumeil et al., 2019; Parks et al., 2020) to the order *Lachnospirales* and to the family *Lachnospiraceae*

**Description of *Candidatus* Physconaster merdicola** sp. nov.

*Candidatus* Physconaster merdicola (mer.di’co.la. L. gen. fem. n. *merda*, faeces; L. masc./fem. suff. *-cola*, inhabitant of; N.L. fem. n. *merdicola* inhanbitant of faeces).

A bacterial species identified by metagenomic analyses. This species includes all bacteria with genomes that show ≥95% average nucleotide identity to the type genome for the species to which we have assigned the genome identifier T0_METABAT__254 and which is available via NCBI BioSample SAMN18871326. This is a new name for the alphanumeric GTDB species sp003507655. The GC content of the type genome is 41.91 % and the genome length is 2.16 Mbp.

**Description of *Candidatus* Ruminococcus anthropi** sp. nov.

*Candidatus* Ruminococcus anthropi (an.thro’pi. Gr. masc. n. *anthropos*, a human being; N.L. gen. masc. n. *anthropi*, of a human being).

A bacterial species identified by metagenomic analyses. This species includes all bacteria with genomes that show ≥95% average nucleotide identity to the type genome for the species to which we have assigned the genome identifier hylon_METABAT__215 and which is available via NCBI BioSample SAMN18871209. This is a new name for the alphanumeric GTDB species sp900314705. GTDB has assigned this species to genus with an alphabetic suffix which cannot be incorporated into a well-formed binomial, so in naming this species, we have used the basonym for the genus. The GC content of the type genome is 33.46 % and the genome length is 1.46 Mbp.

**Description of *Candidatus* Ruminococcus faecihominis** sp. nov.

*Candidatus* Ruminococcus faecihominis (fae.ci.ho’mi.nis. L. fem. n. *faex, faecis* faeces; L. gen. masc. n. *hominis*, of a human being; N.L. gen. n. *faecihominis*, of human faeces).

A bacterial species identified by metagenomic analyses. This species includes all bacteria with genomes that show ≥95% average nucleotide identity to the type genome for the species to which we have assigned the genome identifier T0_METABAT__198 and which is available via NCBI BioSample SAMN18871316. This is a new name for the alphanumeric GTDB species sp003011855. GTDB has assigned this species to genus with an alphabetic suffix which cannot be incorporated into a well-formed binomial, so in naming this species, we have used the basonym for the genus. The GC content of the type genome is 44.64 % and the genome length is 2.86 Mbp.

**Description of *Candidatus* Ruminococcus hominis** sp. nov.

*Candidatus* Ruminococcus hominis (ho’mi.nis. L. gen. masc. n. *hominis*, of a human being).

A bacterial species identified by metagenomic analyses. This species includes all bacteria with genomes that show ≥95% average nucleotide identity to the type genome for the species to which we have assigned the genome identifier rmaize_MAXBIN__013 and which is available via NCBI BioSample SAMN18871266. This is a new name for the alphanumeric GTDB species sp000433635. GTDB has assigned this species to genus with an alphabetic suffix which cannot be incorporated into a well-formed binomial, so in naming this species, we have used the basonym for the genus. The GC content of the type genome is 45.98 % and the genome length is 2.41 Mbp.

**Description of *Candidatus* Sangerella** gen. nov.

*Candidatus* Sangerella (San.ger.el’la. L. fem. dim. suff. *-ella* diminutive ending; N.L. fem. n. *Sangerella* named in honour of Frederick Sanger (1918-2013), British scientist; awarded the 1958 Nobel Prize in Chemistry for his work on the structure of protein and the 1980 Nobel Prize in Chemistry for inventing dideoxy sequencing).

A bacterial genus identified by metagenomic analyses of human faeces. The genus includes all bacteria with genomes that show ≥60% average aminoacid identity to the genome of the type strain from the type species, *Candidatus* Sangerella faecicola. This is a new name for the GTDB alphanumeric genus UBA737. This genus has been assigned by GTDB-Tk v1.5.0 working on GTDB R06-RS202 reference data (Chaumeil et al., 2019; Parks et al., 2020) to the order *Oscillospirales* and to the family *Acutalibacteraceae*

**Description of *Candidatus* Sangerella faecicola** sp. nov.

*Candidatus* Sangerella faecicola (fae.ci’co.la. L. fem. n. *faex, faecis* faeces; L. suff. *- cola* inhabitant of; N.L. fem. n. *faecicola* a microbe inhabitating faeces).

A bacterial species identified by metagenomic analyses. This species includes all bacteria with genomes that show ≥95% average nucleotide identity to the type genome for the species to which we have assigned the genome identifiers rmaize_MAXBIN__077 and T0_METABAT__221 and which is available via NCBI BioSample SAMN18871271. This is a new name for the alphanumeric GTDB species sp900549055. The GC content of the type genome are 47.36 % and 46.01 % and the genome lengths are 2.02 Mbp and 2.89 Mbp.

**Description of *Candidatus* Splanchousia** gen. nov.

*Candidatus* Splanchousia (Splanch.ou’si.a. Gr. neut. n. *splanchnon* guts; L. fem. n. *ousia* an essence; *Splanchousia* a microbe associated with the intestines).

A bacterial genus identified by metagenomic analyses of human faeces. The genus includes all bacteria with genomes that show ≥60% average aminoacid identity to the genome of the type strain from the type species, *Candidatus* Splanchousia colicola. This is a new name for the GTDB alphanumeric genus UBA1191. This genus has been assigned by GTDB-Tk v1.5.0 working on GTDB R06-RS202 reference data (Chaumeil et al., 2019; Parks et al., 2020) to the order *Peptostreptococcales* and to the family *Anaerovoracaceae*

**Description of *Candidatus* Splanchousia colicola** sp. nov.

*Candidatus* Splanchousia colicola (co.li’co.la. L. neut. n. *colum*, colon; L. masc./fem. suff. *-cola*, inhabitant of; N.L. fem. n. *colicola* inhabitant of the colon).

A bacterial species identified by metagenomic analyses. This species includes all bacteria with genomes that show ≥95% average nucleotide identity to the type genome for the species to which we have assigned the genome identifier avicell_METABAT__39 and which is available via NCBI BioSample SAMN18871194. This is a new name for the alphanumeric GTDB species sp900066305. The GC content of the type genome is 49.21 % and the genome length is 2.02 Mbp.

**Description of *Candidatus* Splanchousia faecium** sp. nov.

*Candidatus* Splanchousia faecium (fae’ci.um. L. fem. gen. pl. n. *faecium*, of faeces).

A bacterial species identified by metagenomic analyses. This species includes all bacteria with genomes that show ≥95% average nucleotide identity to the type genome for the species to which we have assigned the genome identifier rmaize_METABAT__166 and which is available via NCBI BioSample SAMN18871279. This is a new name for the alphanumeric GTDB species sp900549125. The GC content of the type genome is 47.53 % and the genome length is 2.11 Mbp.

**Description of *Candidatus* Wallaceimonas** gen. nov.

*Candidatus* Wallaceimonas (Wal.lace.i.mo’nas. L. fem. n. *monas* a monad; N.L. fem. n. *Wallaceimonas* named in honour of British naturalist Alfred Russel Wallace (1823-1913), co-discoverer of evolution by natural selection).

A bacterial genus identified by metagenomic analyses of human faeces. The genus includes all bacteria with genomes that show ≥60% average aminoacid identity to the genome of the type strain from the type species, *Candidatus* Wallaceimonas faecalis. This is a new name for the GTDB alphanumeric genus UMGS1696. This genus has been assigned by GTDB-Tk v1.5.0 working on GTDB R06-RS202 reference data (Chaumeil et al., 2019; Parks et al., 2020) to the order *Oscillospirales* and to the family *CAG-272*

**Description of *Candidatus* Wallaceimonas faecalis** sp. nov.

*Candidatus* Wallaceimonas faecalis (fae.ca’lis. N.L. fem. adj. *faecalis*, of faeces).

A bacterial species identified by metagenomic analyses. This species includes all bacteria with genomes that show ≥95% average nucleotide identity to the type genome for the species to which we have assigned the genome identifier T0_METABAT__60 and which is available via NCBI BioSample SAMN18871330. This is a new name for the alphanumeric GTDB species sp900753285. The GC content of the type genome is 49.14 % and the genome length is 1.81 Mbp.

## Notes

http://www.ncbi.nlm.nih.gov/bioproject/722408

### Competing Interest Statement

The authors have declared no competing interest.

### Summary of Updates

Table 1 (species protologues names) added, and updated on Supp table 7. Some referencing errors corrected

## References

1. Kau AL, Ahern PP, Griffin NW, Goodman AL, Gordon JI. Human nutrition, the gut microbiome and the immune system. Nature. 2011;474(7351):327–36.

2. Koropatkin NM, Cameron EA, Martens EC. How glycan metabolism shapes the human gut microbiota. Nature Reviews Microbiology. 2012;10(5):323–35.

3. Chambers ES, Byrne CS, Morrison DJ, Murphy KG, Preston T, Tedford C, et al. Dietary supplementation with inulin-propionate ester or inulin improves insulin sensitivity in adults with overweight and obesity with distinct effects on the gut microbiota, plasma metabolome and systemic inflammatory responses: a randomised cross-over trial. Gut. 2019;68(8):1430–8.

4. Blaak E, Canfora E, Theis S, Frost G, Groen A, Mithieux G, et al. Short chain fatty acids in human gut and metabolic health. Beneficial Microbes. 2020;11(5):411–55.

5. Lloyd-Price J, Mahurkar A, Rahnavard G, Crabtree J, Orvis J, Hall AB, et al. Strains, functions and dynamics in the expanded Human Microbiome Project. Nature. 2017;550(7674):61–6.

6. Martens EC, Kelly AG, Tauzin AS, Brumer H. The devil lies in the details: how variations in polysaccharide fine-structure impact the physiology and evolution of gut microbes. Journal of molecular biology. 2014;426(23):3851–65.

7. Warren FJ, Fukuma NM, Mikkelsen D, Flanagan BM, Williams BA, Lisle AT, et al. Food starch structure impacts gut microbiome composition. Msphere. 2018;3(3).

8. Deehan EC, Yang C, Perez-Muñoz ME, Nguyen NK, Cheng CC, Triador L, et al. Precision microbiome modulation with discrete dietary fiber structures directs short-chain fatty acid production. Cell Host & Microbe. 2020.

9. Carmody RN, Bisanz JE, Bowen BP, Maurice CF, Lyalina S, Louie KB, et al. Cooking shapes the structure and function of the gut microbiome. Nature microbiology. 2019;4(12):2052–63.

10. Lapébie P, Lombard V, Drula E, Terrapon N, Henrissat B. Bacteroidetes use thousands of enzyme combinations to break down glycans. Nature communications. 2019;10(1):1–7.

11. Kujawska M, La Rosa SL, Pope PB, Hoyles L, McCartney AL, Hall LJ. Succession of Bifidobacterium longum strains in response to the changing early-life nutritional environment reveals specific adaptations to distinct dietary substrates. 2020.

12. Charalampous T, Kay GL, Richardson H, Aydin A, Baldan R, Jeanes C, et al. Nanopore metagenomics enables rapid clinical diagnosis of bacterial lower respiratory infection. Nature Biotechnology. 2019;37(7):783–92.

13. De Coster W, De Rijk P, De Roeck A, De Pooter T, D’Hert S, Strazisar M, et al. Structural variants identified by Oxford Nanopore PromethION sequencing of the human genome. Genome research. 2019;29(7):1178–87.

14. Bertrand D, Shaw J, Kalathiyappan M, Ng AHQ, Kumar MS, Li C, et al. Hybrid metagenomic assembly enables high-resolution analysis of resistance determinants and mobile elements in human microbiomes. Nature biotechnology. 2019;37(8):937–44.

15. Arumugam K, Bağcı C, Bessarab I, Beier S, Buchfink B, Gorska A, et al. Annotated bacterial chromosomes from frame-shift-corrected long-read metagenomic data. Microbiome. 2019;7(1):61.

16. Singleton CM, Petriglieri F, Kristensen JM, Kirkegaard RH, Michaelsen TY, Andersen MH, et al. Connecting structure to function with the recovery of over 1000 high-quality activated sludge metagenome-assembled genomes encoding full-length rRNA genes using long-read sequencing. bioRxiv. 2020.

17. Stewart RD, Auffret MD, Warr A, Walker AW, Roehe R, Watson M. Compendium of 4,941 rumen metagenome-assembled genomes for rumen microbiome biology and enzyme discovery. Nature biotechnology. 2019;37(8):953.

18. Walker AW, Duncan SH, Leitch ECM, Child MW, Flint HJ. pH and peptide supply can radically alter bacterial populations and short-chain fatty acid ratios within microbial communities from the human colon. Appl Environ Microbiol. 2005;71(7):3692–700.

19. Leitch ECM, Walker AW, Duncan SH, Holtrop G, Flint HJ. Selective colonization of insoluble substrates by human faecal bacteria. Environmental microbiology. 2007;9(3):667–79.

20. Bowers RM, Kyrpides NC, Stepanauskas R, Harmon-Smith M, Doud D, Reddy T, et al. Minimum information about a single amplified genome (MISAG) and a metagenome-assembled genome (MIMAG) of bacteria and archaea. Nature biotechnology. 2017;35(8):725–31.

21. Pallen MJ, Telatin A, Oren A. The Next Million Names for Archaea and Bacteria. Trends Microbiol. 2020.

22. Machovič M, Janeček Š. Domain evolution in the GH13 pullulanase subfamily with focus on the carbohydrate-binding module family 48. Biologia. 2008;63(6):1057–68.

23. Moss EL, Maghini DG, Bhatt AS. Complete, closed bacterial genomes from microbiomes using nanopore sequencing. Nature Biotechnology. 2020:1–7.

24. Maghini DG, Moss EL, Vance SE, Bhatt AS. Improved high-molecular-weight DNA extraction, nanopore sequencing and metagenomic assembly from the human gut microbiome. Nature Protocols. 2021;16(1):458–71.

25. Aagaard K, Petrosino J, Keitel W, Watson M, Katancik J, Garcia N, et al. The Human Microbiome Project strategy for comprehensive sampling of the human microbiome and why it matters. The FASEB Journal. 2013;27(3):1012–22.

26. Methé BA, Nelson KE, Pop M, Creasy HH, Giglio MG, Huttenhower C, et al. A framework for human microbiome research. Nature. 2012;486(7402):215.

27. Campbell JM, Fahey Jr GC, Wolf BW. Selected indigestible oligosaccharides affect large bowel mass, cecal and fecal short-chain fatty acids, pH and microflora in rats. The Journal of nutrition. 1997;127(1):130–6.

28. Lombard V, Golaconda Ramulu H, Drula E, Coutinho PM, Henrissat B. The carbohydrate-active enzymes database (CAZy) in 2013. Nucleic acids research. 2014;42(D1):D490–D5.

29. El Kaoutari A, Armougom F, Gordon JI, Raoult D, Henrissat B. The abundance and variety of carbohydrate-active enzymes in the human gut microbiota. Nature Reviews Microbiology. 2013;11(7):497–504.

30. Benítez-Páez A, Gómez del Pulgar EM, Sanz Y. The glycolytic versatility of Bacteroides uniformis CECT 7771 and its genome response to oligo and polysaccharides. Frontiers in cellular and infection microbiology. 2017;7:383.

31. Moens F, Weckx S, De Vuyst L. Bifidobacterial inulin-type fructan degradation capacity determines cross-feeding interactions between bifidobacteria and Faecalibacterium prausnitzii. International journal of food microbiology. 2016;231:76–85.

32. Ramirez-Farias C, Slezak K, Fuller Z, Duncan A, Holtrop G, Louis P. Effect of inulin on the human gut microbiota: stimulation of Bifidobacterium adolescentis and Faecalibacterium prausnitzii. British Journal of Nutrition. 2008;101(4):541–50.

33. Flint HJ, Scott KP, Duncan SH, Louis P, Forano E. Microbial degradation of complex carbohydrates in the gut. Gut microbes. 2012;3(4):289–306.

34. Bui TPN, Schols HA, Jonathan M, Stams AJ, de Vos WM, Plugge CM. Mutual Metabolic Interactions in Co-cultures of the Intestinal Anaerostipes rhamnosivorans With an Acetogen, Methanogen, or Pectin-Degrader Affecting Butyrate Production. Frontiers in microbiology. 2019;10:2449.

35. Ze X, David YB, Laverde-Gomez JA, Dassa B, Sheridan PO, Duncan SH, et al. Unique organization of extracellular amylases into amylosomes in the resistant starch-utilizing human colonic Firmicutes bacterium Ruminococcus bromii. MBio. 2015;6(5).

36. Upadhyaya B, McCormack L, Fardin-Kia AR, Juenemann R, Nichenametla S, Clapper J, et al. Impact of dietary resistant starch type 4 on human gut microbiota and immunometabolic functions. Scientific reports. 2016;6:28797.

37. Xie Z, Wang S, Wang Z, Fu X, Huang Q, Yuan Y, et al. In vitro fecal fermentation of propionylated high-amylose maize starch and its impact on gut microbiota. Carbohydrate polymers. 2019;223:115069.

38. Ryan SM, Fitzgerald GF, van Sinderen D. Screening for and identification of starch-, amylopectin-, and pullulan-degrading activities in bifidobacterial strains. Applied and Environmental Microbiology. 2006;72(8):5289–96.

39. Crittenden R, Laitila A, Forssell P, Mättö J, Saarela M, Mattila-Sandholm T, et al. Adhesion of bifidobacteria to granular starch and its implications in probiotic technologies. Applied and Environmental Microbiology. 2001;67(8):3469–75.

40. Williams BA, Bosch MW, Boer H, Verstegen MW, Tamminga S. An in vitro batch culture method to assess potential fermentability of feed ingredients for monogastric diets. Animal Feed Science and Technology. 2005;123:445–62.

41. De Coster W, D’Hert S, Schultz DT, Cruts M, Van Broeckhoven C. NanoPack: visualizing and processing long-read sequencing data. Bioinformatics. 2018;34(15):2666–9; doi: 10.1093/bioinformatics/bty149.

42. Chen S, Zhou Y, Chen Y, Gu J. fastp: an ultra-fast all-in-one FASTQ preprocessor. Bioinformatics. 2018;34(17):i884–i90.

43. Truong DT, Franzosa EA, Tickle TL, Scholz M, Weingart G, Pasolli E, et al. MetaPhlAn2 for enhanced metagenomic taxonomic profiling. Nature methods. 2015;12(10):902–3.

44. Li D, Liu C-M, Luo R, Sadakane K, Lam T-W. MEGAHIT: an ultra-fast single-node solution for large and complex metagenomics assembly via succinct de Bruijn graph. Bioinformatics. 2015;31(10):1674–6.

45. Li D, Luo R, Liu C-M, Leung C-M, Ting H-F, Sadakane K, et al. MEGAHIT v1. 0: A fast and scalable metagenome assembler driven by advanced methodologies and community practices. Methods. 2016;102:3–11.

46. Bertrand D, Shaw J, Kalathiyappan M, Ng AHQ, Kumar MS, Li C, et al. Hybrid metagenomic assembly enables high-resolution analysis of resistance determinants and mobile elements in human microbiomes. Nat Biotechnol. 2019;37(8):937–44; doi: 10.1038/s41587-019-0191-2.

47. Wu Y-W, Simmons BA, Singer SW. MaxBin 2.0: an automated binning algorithm to recover genomes from multiple metagenomic datasets. Bioinformatics. 2016;32(4):605–7.

48. Kang DD, Froula J, Egan R, Wang Z. MetaBAT, an efficient tool for accurately reconstructing single genomes from complex microbial communities. PeerJ. 2015;3:e1165; doi: 10.7717/peerj.1165.

49. Eren AM, Kiefl E, Shaiber A, Veseli I, Miller SE, Schechter MS, et al. Community-led, integrated, reproducible multi-omics with anvi’o. Nature Microbiology. 2021;6(1):3–6.

50. Sieber CMK, Probst AJ, Sharrar A, Thomas BC, Hess M, Tringe SG, et al. Recovery of genomes from metagenomes via a dereplication, aggregation and scoring strategy. Nat Microbiol. 2018;3(7):836–43; doi: 10.1038/s41564-018-0171-1.

51. Parks DH, Imelfort M, Skennerton CT, Hugenholtz P, Tyson GW. CheckM: assessing the quality of microbial genomes recovered from isolates, single cells, and metagenomes. Genome Res. 2015;25(7):1043–55; doi: 10.1101/gr.186072.114.

52. Olm MR, Brown CT, Brooks B, Banfield JF. dRep: a tool for fast and accurate genomic comparisons that enables improved genome recovery from metagenomes through de-replication. The ISME journal. 2017;11(12):2864–8.

53. Zhang H, Yohe T, Huang L, Entwistle S, Wu P, Yang Z, et al. dbCAN2: a meta server for automated carbohydrate-active enzyme annotation. Nucleic Acids Research. 2018;46(W1):W95–W101.

54. Finn RD, Clements J, Eddy SR. HMMER web server: interactive sequence similarity searching. Nucleic acids research. 2011;39(suppl_2):W29–W37.

55. Buchfink B, Xie C, Huson DH. Fast and sensitive protein alignment using DIAMOND. Nature methods. 2015;12(1):59–60.

56. Busk PK, Pilgaard B, Lezyk MJ, Meyer AS, Lange L. Homology to peptide pattern for annotation of carbohydrate-active enzymes and prediction of function. BMC bioinformatics. 2017;18(1):214.

