## Supplementary figures and images for "Linking carbohydrate structure with function in the human gut microbiome using hybrid metagenome assemblies"

### Supplemental Figure 1

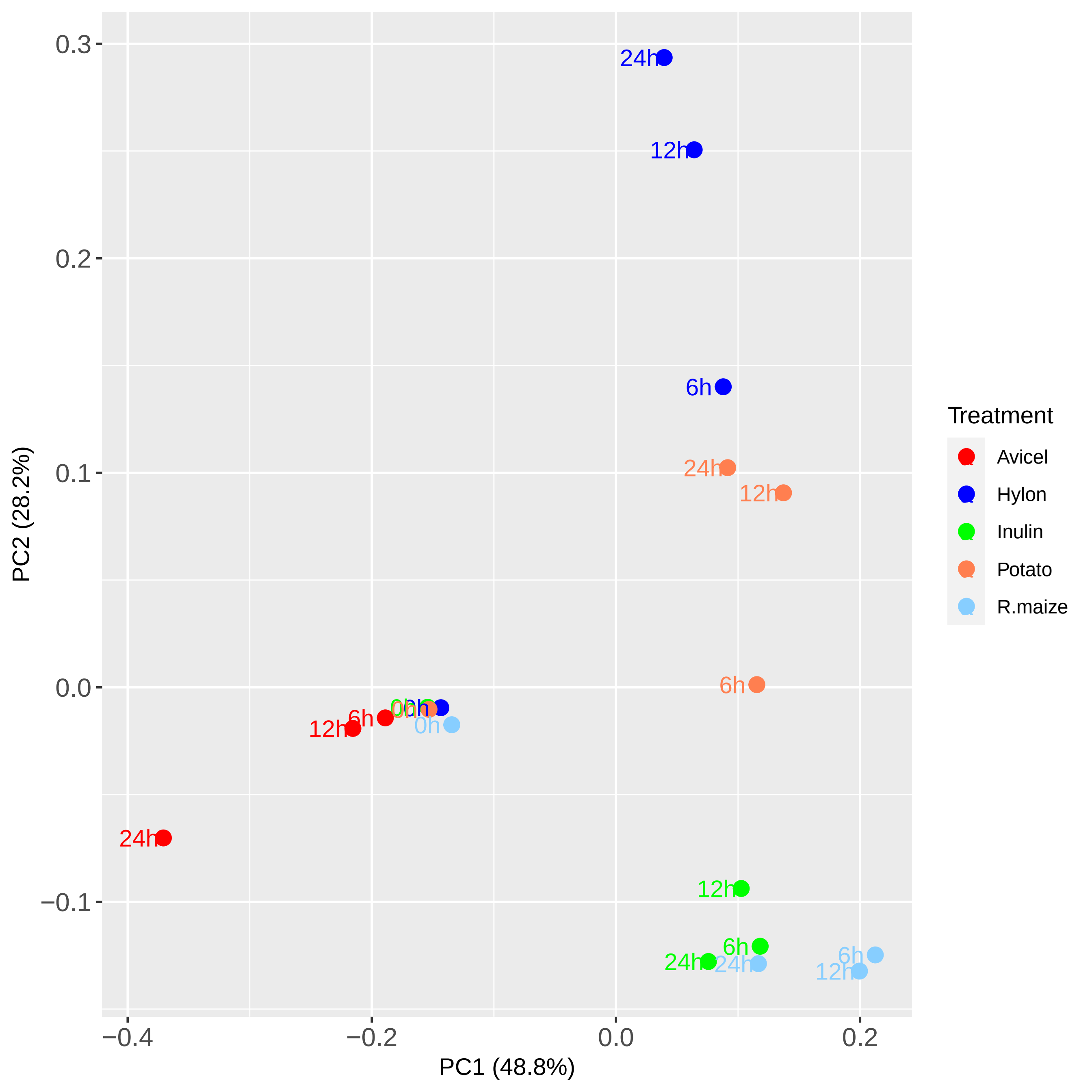

### Supplemental Figure 2

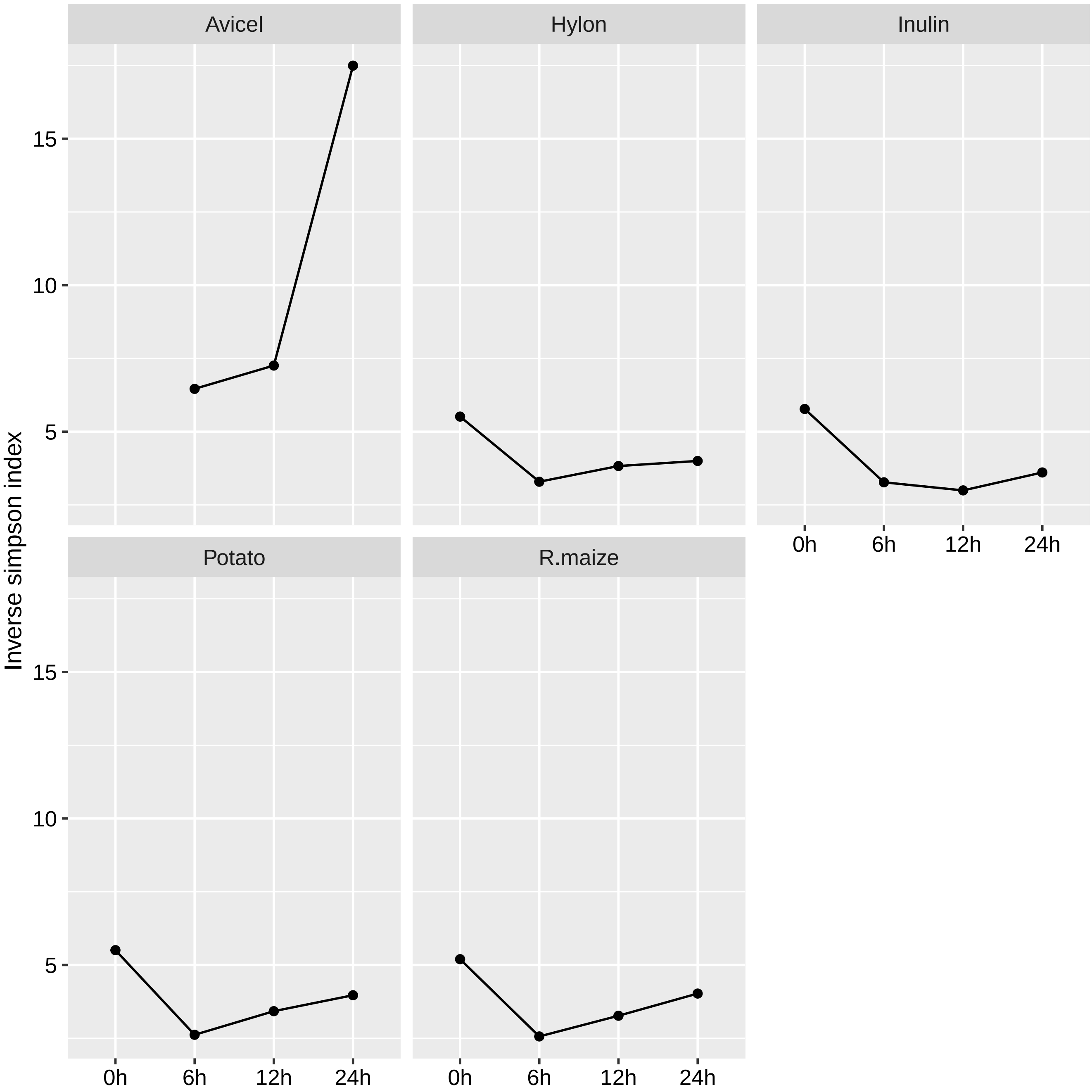

### Supplemental Figure 3

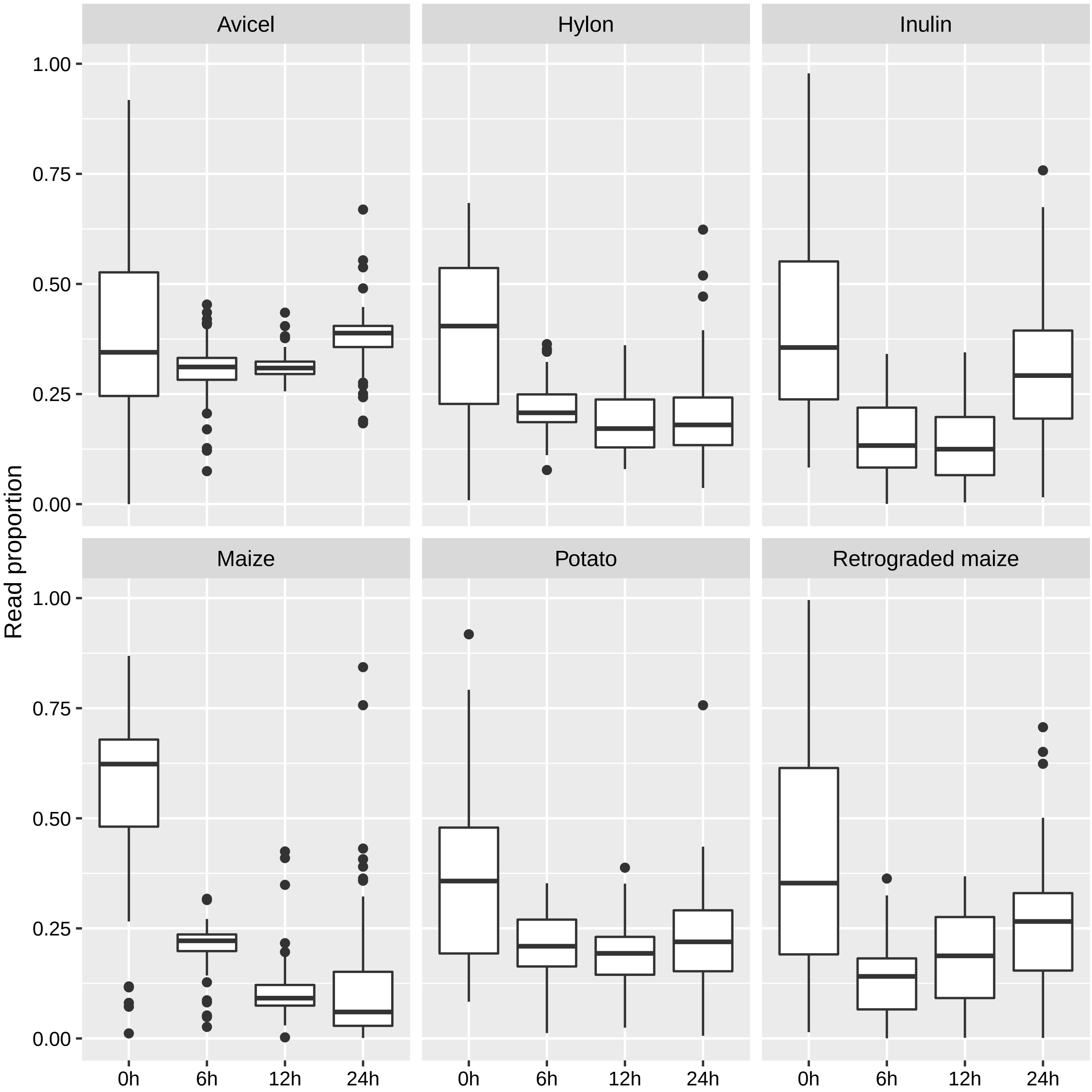

### Supplemental Figure 4

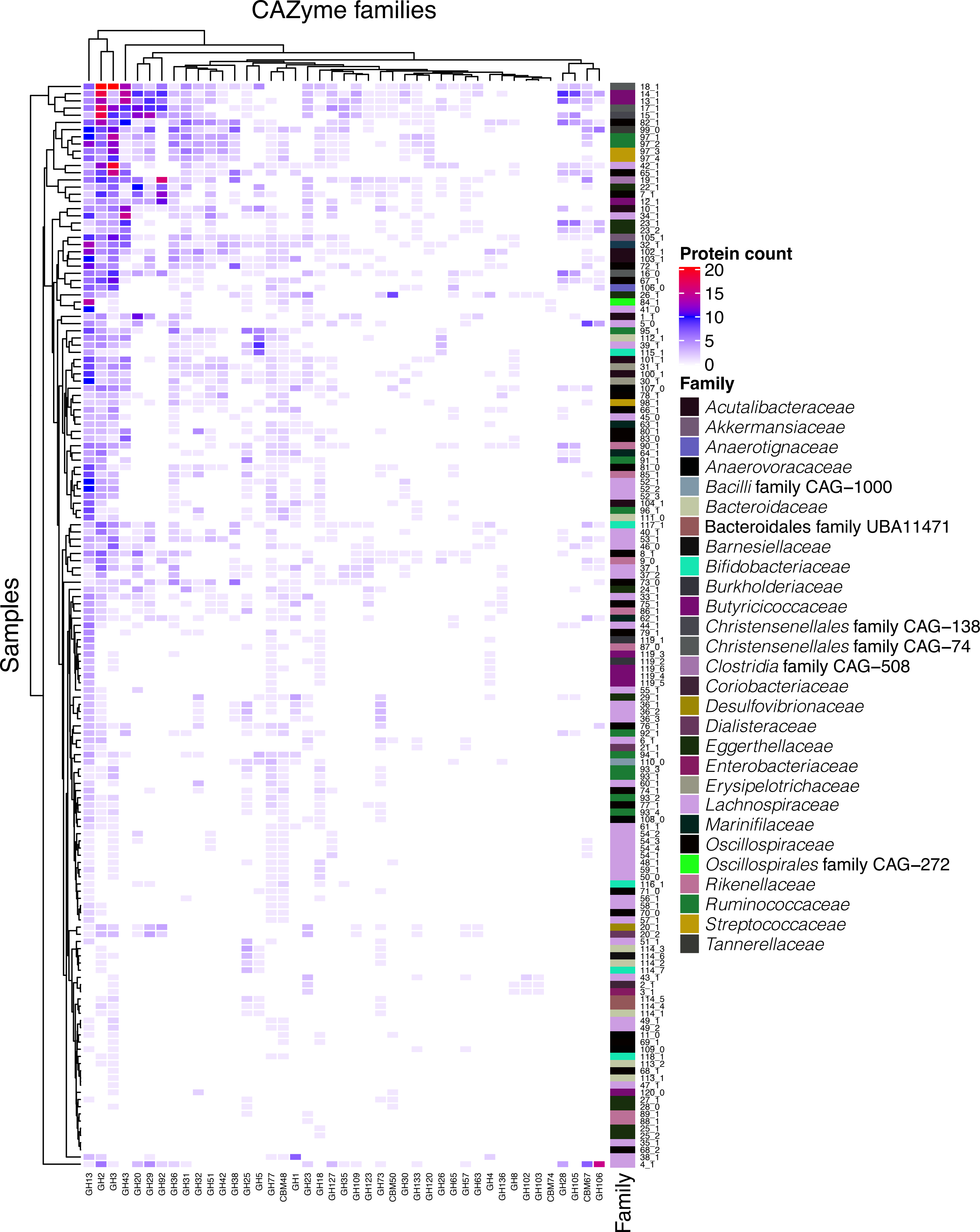
