## Supplemental Table 3 for "Linking carbohydrate structure with function in the human gut microbiome using hybrid metagenome assemblies"

Supplementary Table 3: Assembly statistics

| Treatment | Number of contigs | N50 (Kbp) | Largest Contig | Total assembly length |
| --- | --- | --- | --- | --- |
| short read assembly (using Megahit) | | | | |
| Avicell | 362,193 | 5.9 | 522,610 | 611,883,067 |
| Hylon | 359,933 | 4.9 | 807,467 | 566,999,279 |
| Inulin | 325,487 | 4.5 | 667,421 | 478,846,585 |
| Potato | 331,720 | 5.7 | 580,261 | 539,511,443 |
| R.maize | 317,785 | 4.7 | 703,349 | 487,894,115 |
| N.Maize | 304,686 | 5.2 | 522,608 | 494,497,388 |
| Time 0h | 369,675 | 6.2 | 551,334 | 646,904,987 |
| Hybrid assembly (using OPERA-MS with Megahit and minimap2) | | | | |
| Avicell | 336,341 | 10.0 | 1,204,096 | 619,505,714 |
| Hylon | 268,928 | 29.2 | 1,635,802 | 615,331,172 |
| Inulin | 236,720 | 36.0 | 1,544,219 | 532,364,427 |
| Potato | 303,698 | 10.8 | 1,632,261 | 550,284,682 |
| R.maize | 226,040 | 38.1 | 1,148,593 | 540,634,515 |
| N.Maize | 288,406 | 7.7 | 1,052,519 | 499,371,327 |
| Time 0h | 278,441 | 33.5 | 1,381,878 | 694,064,798 |
