## Supplemental Table 6 for "Linking carbohydrate structure with function in the human gut microbiome using hybrid metagenome assemblies"

Supplementary Table 6: GTDb taxonomy from the dereplication MAG clusters

| Phylum | Family | Dereplicated genomes | No. of genus | No. of species | No. of MAGs |
| --- | --- | --- | --- | --- | --- |
| *Actinobacteriota* | *Bifidobacteriaceae* | 5 | 1 | 5 | 29 |
|  | *Coriobacteriaceae* | 1 | 1 | 1 | 7 |
|  | *Eggerthellaceae* | 10 | 3 | 8 | 16 |
| *Bacteroidota* | *Bacteroidaceae* | 7 | 4 | 6 | 28 |
|  | *Barnesiellaceae* | 1 | 1 | 1 | 7 |
|  | *Marinifilaceae* | 3 | 2 | 2 | 6 |
|  | *Rikenellaceae* | 7 | 3 | 6 | 21 |
|  | *Tannerellaceae* | 1 | 1 | 1 | 6 |
|  | *UBA11471* | 2 | 1 | 1 | 4 |
| *Desulfobacterota_A* | *Desulfovibrionaceae* | 1 | 1 | 1 | 7 |
| *Firmicutes* | *CAG-1000* | 1 | 1 | 1 | 1 |
|  | *Erysipelotrichaceae* | 2 | 2 | 2 | 13 |
|  | *Streptococcaceae* | 3 | 1 | 1 | 5 |
|  | UBA660 | 1 | 1 | 1 | 1 |
| *Firmicutes_A* | *Acutalibacteraceae* | 8 | 7 | 7 | 31 |
|  | *Anaerotignaceae* | 1 | 1 | 1 | 1 |
|  | *Anaerovoracaceae* | 4 | 2 | 2 | 7 |
|  | *Butyricicoccaceae* | 8 | 2 | 2 | 12 |
|  | *CAG-138* | 1 | 1 | 1 | 1 |
|  | *CAG-272* | 1 | 1 | 1 | 1 |
|  | *CAG-508* | 1 | 1 | 1 | 1 |
|  | *CAG-74* | 3 | 2 | 2 | 15 |
|  | *Lachnospiraceae* | 41 | 30 | 35 | 130 |
|  | *Oscillospiraceae* | 22 | 7 | 9 | 79 |
|  | *Ruminococcaceae* | 11 | 7 | 9 | 50 |
| *Firmicutes_C* | *Dialisteraceae* | 2 | 2 | 2 | 8 |
| *Proteobacteria* | *Burkholderiaceae* | 2 | 2 | 2 | 13 |
|  | *Enterobacteriaceae* | 1 | 1 | 1 | 4 |
| *Verrucomicrobiota* | *Akkermansiaceae* | 1 | 1 | 1 | 7 |
