## Supplemental Table 10 for "Linking carbohydrate structure with function in the human gut microbiome using hybrid metagenome assemblies"

| Treatment | Early degraders | | Late degraders | |
| --- | --- | --- | --- | --- |
|  | Cluster | Taxa (ordered by cluster) | Cluster | Taxa (ordered by cluster) |
| Avicell | cluster_111_0,  cluster_26_1,  cluster_28_0 | *Blautia hydrogenotrophica, Escherichia coli, Candidatus* Splanchousia colicola | cluster_51_1,  cluster_58_1,  cluster_63_1 | *Candidatus Caccadaptatus darwinii*, *Candidatus Minthonaster hominis*, *Faecalibacterium prausnitzii* |
| Hylon | cluster_104_1,  cluster_26_1,  cluster_32_1,  cluster_41_0,  cluster_84_1  cluster_65_1,  cluster_72_1 | *Pararoseburia caccae*  , *Escherichia coli, Bifidobacterium adolescentis, Candidatus* Ruminococcus anthropi*,Ruminococcus bromii, Gemmiger qucibialis, Candidatus* Eisenbergiella faecalis | cluster_49_1,  cluster_51_1,  cluster_52_1,  cluster_58_1,  cluster_96_1 | *Dysosmobacter segnis*, *Candidatus Caccadaptatus darwinii*, *Candidatus Enteromorpha quadrami*, *Candidatus Minthonaster hominis*,  *Candidatus Blautia hennigii* |
| Inulin | cluster_29_1,  cluster_38_1,  cluster_82_1 | *Candidatus* Colinsella sterocoris*,*  *Candidatus* Holdemanella enterica, *Candidatus* Minthovivens enterohominis | cluster_18_1  cluster_4_1  cluster_49_2  cluster_51_1  cluster_56_1  cluster_58_1  cluster_63_1 | *Bacteroides uniformis, Alistipes indistinctus,* *Dysosmobacter segnis*, *Candidatus* Caccadaptatus darwinii, *Candidatus* Minthonaster faecium, *Candidatus* Minthonaster hominis*, Faecalibacterium prausnitzii* |
| Potato | cluster_2_1  cluster_29_1  cluster_44_1  cluster_63_1  cluster_66_1  cluster_84_1  cluster_96_1 | *Sutterella wadsworthensis,* *Candidatus* Colinsella sterocoris*,* *Candidatus* *Aphodonaster merdae*,  *Faecalibacterium prausnitzii, Candidatus* Gemmiger merdicola*, Ruminococcus bromii, Candidatus* Blautia hennigii | cluster_8_1  cluster_48_1  cluster_51_1  cluster_62_1 | *Alistipes shahii,* *Dysosmobacter welbionis, Candidatus* Caccadaptatus darwinii,  *Candidatus* Colihabitans norwichensis |
| R.maize | cluster_17_1  cluster_19_1  cluster_2_1  cluster_27_1  cluster_4_1  cluster_43_1  cluster_48_1  cluster_51_1  cluster_58_1  cluster_87_0 | *Bacteroides fragilis, Parabacteroides* diastonis, *Sutterella wadsworthensis, Candidatus* Splanchousia faecium*, Alistipes indistinctus, Bilophila wadsworthia,*  *Dysosmobacter welbionis, Candidatus* Caccadaptatus darwinii, *Candidatus* Minthonaster hominis, *Candidatus* Minthomorpha faecalis | cluster_26_1  cluster_29_1  cluster_30_1  cluster_31_1  cluster_32_1  cluster_34_1  cluster_38_1  cluster_46_0 | *Escherichia coli,* *Candidatus* Colinsella sterocoris*, Bifidobacterium animalis, Bifidobacterium catenulatum, Bifidobacterium adolescentis, Bifidobactrium longum, Candidatus* Holdemanella enterica*, Ruthenibacterium lactatiformans* |

Supplementary table 10: Genomes depicted as early and late degraders according to the time the genomes showed a 2x fold change.
