## Supplemental Table 13 for "Linking carbohydrate structure with function in the human gut microbiome using hybrid metagenome assemblies"

| **Supplementary Table 13** Media preparation materials, sources, and quantity^a^ | | |
| --- | --- | --- |
| **Solution** | **Source** | **Quantity/litre** |
| ***Fatty Acid Solution*** | | |
| NaOH (0.2M) | Sigma-Aldrich, catalogue no 06203 | 1 L |
| Acetic acid | Sigma-Aldrich, catalogue no A6283 | 6.85 mL |
| Propionic acid | Sigma-Aldrich, catalogue no 402907 | 3.00 mL |
| Butyric acid | Sigma-Aldrich, catalogue no B103500 | 1.84 mL |
| Isobutyric acid | Sigma-Aldrich, catalogue no 58360 | 0.47 mL |
| 2-Methylbutyric acid | Sigma-Aldrich, catalogue no W269514 | 0.55 mL |
| Valeric Acid | Sigma-Aldrich, catalogue no 240370 | 0.55 mL |
| Isovaleric acid | Sigma-Aldrich, catalogue no 129542 | 0.55 mL |
| ***Haemin Solution*** | | |
| NaOH (0.05M) | Sigma-Aldrich, catalogue no 06203 | 1 L |
| Haemin |  | 100.0 mg |
| ***Trace Mineral Solution*** | | |
| HCl (0.02M) | Fisher Scientific, catalogue no 15676840 | 1 L |
| Manganese chloride (MnCl_2_•4H_2_O) | Sigma-Aldrich, catalogue no M3634 | 25.0 mg |
| Ferrous Sulphate (FeSO_4_•7H_2_O) | Sigma-Aldrich, catalogue no F7002 | 20.0 mg |
| Zinc chloride (ZnCl_2_) | Sigma-Aldrich, catalogue no 793523 | 25.0 mg |
| Copper chloride (CuCl•2H_2_O) | Sigma-Aldrich, catalogue no 307483 | 25.0 mg |
| Cobalt chloride (CoCl_2_•6H_2_O) | Sigma-Aldrich, catalogue no 255599 | 50.0 mg |
| Selenium dioxide (SeO_2_) | Sigma-Aldrich, catalogue no 213365 | 50.0 mg |
| Nickel chloride (NiCl_2_•6H_2_O) | Sigma-Aldrich, catalogue no 223387 | 250.0 mg |
| Sodium molybdate (Na_2_MoO_4_•2H_2_O) | Sigma-Aldrich, catalogue no 331058 | 250.0 mg |
| Sodium metavanadate (NaVO_3_) | Sigma-Aldrich, catalogue no 590088 | 31.4 mg |
| Boric acid (H_3_BO_3_) | Sigma-Aldrich, catalogue no 31146 | 250.0 mg |
| ***Vitamin-Phosphate Solution*** (filtered sterilized using 0.2μm nylon filter) | | |
| Potassium phosphate monobasic KH_2_PO_4_ | Sigma-Aldrich, catalogue no P9791 | 20.4 mg |
| Biotin | Sigma-Aldrich, catalogue no B4639 | 20.6 mg |
| Folic acid | Sigma-Aldrich, catalogue no F8758 | 164.0 mg |
| Calcium D-pantothenate | Sigma-Aldrich, catalogue no P5155 | 164.0 mg |
| Nicotinamide | Sigma-Aldrich, catalogue no 72340 | 164.0 mg |
| Riboflavin | Sigma-Aldrich, catalogue no R9504 | 164.0 mg |
| Thiamine HCl | Sigma-Aldrich, catalogue no T4625 | 164.0 mg |
| Pyridoxine HCl | Sigma-Aldrich, catalogue no 181986 | 164.0 mg |
| *Para*-amino benzoic acid | Sigma-Aldrich, catalogue no A9878 | 20.4 mg |
| Cyanocobalamin (Vitamin B12) | Sigma-Aldrich, catalogue no C3607 | 20.6 mg |
| ***Reducing Agent*** (filtered sterilized using 0.2μm nylon filter) | | |
| Deionized water at 100°C* |  | 1L |
| L-Cysteine HCl | VWR, catalogue no ACRO434850010 | 20.0 g |
| Na_2_S•9H_2_O | Sigma-Aldrich, catalogue no 431648 | 20.0 g |
| ***Sodium Carbonate Solution*** (Na_2_CO_3_, degassed with CO2 and autoclaved) Sigma 223484 | | 82g/L water* |
| ***Vitamin Phosphate+ Na_2_CO_3_ Solution*** | | **75 mL total** |
| Na_2_CO_3_ |  | 60 mL |
| Vitamin Phosphate |  | 15 mL |
| ***Resazurin Solution*** | Sigma-Aldrich, catalogue no R7017 | 1.0 g/L water* |
| ***Potassium Hydroxide*** (3M) | Sigma-Aldrich, catalogue no 221473 | 160 g/L water* |
| ***Basal Solution*** (degassed with CO_2_ and autoclaved) | | |
| Deionized water at 100°C* |  | 1L |
| KCl | Sigma-Aldrich, catalogue no P3911 | 713.4 mg |
| NaCl | Sigma-Aldrich, catalogue no S7653 | 713.4 mg |
| CaCl_2_•2H_2_O | Sigma-Aldrich, catalogue no 223506 | 237.8 mg |
| MgSO_4_•7H_2_O | Sigma-Aldrich, catalogue no | 594.5 mg |
| Pipes buffer | Sigma-Aldrich, catalogue no P6757 | 1,783.5 mg |
| NH_4_Cl | Sigma-Aldrich, catalogue no A9434 | 642.0 mg |
| Trypticase Peptone | VWR, catalogue no 1.07213.1000 | 1,189.0 mg |
| **Reagents** |  |  |
| Resazurin solution |  | 1.17 mL |
| Trace Mineral solution |  | 11.89 mL |
| Haemin solution |  | 11.89 mL |
| Fatty Acid solution |  | 11.89 mL |
| KOH |  | Adjust pH 6.8 |
